# Neoplastic and immune single cell transcriptomics define subgroup-specific intra-tumoral heterogeneity of childhood medulloblastoma

**DOI:** 10.1101/2020.08.28.272021

**Authors:** Kent A. Riemondy, Sujatha Venkataraman, Nicholas Willard, Anandani Nellan, Bridget Sanford, Andrea M. Griesinger, Vladimir Amani, Siddhartha Mitra, Todd C. Hankinson, Michael H. Handler, Martin Sill, Jennifer Ocasio, Seth J. Weir, Daniel S Malawsky, Timothy R. Gershon, Alexandra Garancher, Robert J. Wechsler-Reya, Jay R. Hesselberth, Nicholas K. Foreman, Andrew M. Donson, Rajeev Vibhakar

## Abstract

Medulloblastoma (MB) is a heterogeneous disease in which neoplastic cells and associated immune cells contribute to disease progression. To better understand cellular heterogeneity in MB we use single-cell RNA sequencing, immunohistochemistry and deconvolution of transcriptomic data to profile neoplastic and immune populations in childhood MB samples. Neoplastic cells cluster primarily according to individual sample of origin which is in part due to the effect of chromosomal copy number gains and losses. Harmony alignment reveals novel MB subgroup/subtype-associated subpopulations that recapitulate neurodevelopmental processes and are associated with clinical outcomes, including discrete photoreceptor-like cells in MB subgroups GP3 and GP4 and nodule-associated neuronally-differentiated cells in subgroup SHH. We definitively chart the spectrum of MB immune cell infiltrates, which include subgroup/subtype-associated developmentally-related neuron-pruning and antigen presenting myeloid cells. MB cellular diversity is recapitulated in genetically engineered mouse subgroup-specific models of MB. These findings provide a clearer understanding of both the neoplastic and immune cell heterogeneity in MB.

## Introduction

Medulloblastoma (MB) is an aggressive brain tumor arising predominantly in childhood, comprised of 4 consensus molecular subgroups – GP3, GP4, SHH and WNT – that have been defined through bulk-tumor transcriptomic, methylomic and genomic profiling studies (1). GP3 subgroup tumors often exhibit amplification of *MYC* and are associated with the poorest clinical outcome; SHH and WNT often harbor mutations in sonic hedgehog and wingless pathway genes, respectively; and the genetic driver for GP4 is less clear. Subsequent studies have subdivided MB subgroups into finer subtypes with disparate outcomes and molecular features (2–4). Unfortunately, intra-tumoral cellular heterogeneity inherent in bulk-tumor samples impedes a clearer understanding of MB biology. Neoplastic and immune heterogeneity underlies a tumor’s ability to proliferate, survive and evade therapeutic interventions. The emergence of single cell analysis now allows us to examine the cellular diversity inherent in biological systems, resulting in transformative findings when applied to pediatric brain tumors including MB. Hovestadt *et al.* used SMART-seq single cell RNA-seq (scRNA-seq) to interrogate MB, identifying cellular diversity within individual samples recapitulating cell cycle, progenitor and differentiated neuronal programs (5). Vladoiu *et al.* used droplet-based scRNA-seq to map mouse cerebellar developmental cellular lineages, and then compared these to childhood cerebellar brain tumors including MB samples, identifying putative subgroup-specific cells of origin (6). In the present study, we add significant new knowledge by increasing the number of samples and cells per tumor and by adding MB genetrically-engineered mouse models (GEMMs) for cross-species analysis. .

Our studies identify and characterize more granular differentiation trajectories in detail, especially previously unnoted discrete photoreceptor and glutamatergic subpopulations across GP3 and GP4 tumors. Importantly we describe for the first time considerable immune cell diversity in MB in detail, including subgroup-specific immune populations. In preclinical models, we identify neoplastic subgroup-specific genetically-engineered mouse model (GEMM) subpopulations that correspond to those seen in human samples. Finally, we provide these data in the form of an interactive browser for users to interrogate MB subgroup neoplastic and immune cells and GEMMs at single cell resolution.

## Results

### Sample-specific clustering in MB scRNA-seq is an effect of chromosomal copy number gains and losses

To better understand cellular heterogeneity in MB we performed scRNA-seq on 28 primary childhood MB patient samples and a single matched sample from recurrence. MB samples were classified into subgroups (1 WNT, 9 SHH, 7 GP3 and 11 GP4) based on bulk-tumor methylome and pooled single cell transcriptome profiles (Supplementary Table 1). The SHH, GP3 and GP4 samples were further assigned to more recently described subtypes (2, 4). Using the 10x Genomics droplet-sequencing-based platform we generated transcriptomes at the single-cell level as previously described (7). After excluding poor quality cells based on low unique molecular identifier (UMI) counts and high mitochondrial proportions, 39,946 cells (~1,300 per sample) remained that passed quality controls (Supplementary Fig. 1).

Cell by gene expression matrices projected as 2D UMAP plots revealed multiple clusters of cells that were either unique to or shared between samples. Cell type analysis based on transcriptomic signatures identified 3 main categories of cell types – neoplastic, immune (myeloid and lymphocyte lineages) and non-neoplastic stroma (astrocytes, oligodendrocytes and vascular endothelium) (Fig. 1a). The distinction between neoplastic and non-neoplastic designation was confirmed by examining copy number variants (CNVs) inferred from the single cell RNA-seq data (inferCNV)(Fig 1b). We identified numerous MB subgroup specific copy number gains and losses that were highly concordant with CNVs identified by methylation analysis of matched samples, the most common of these being isochromosome 17q in the majority of GP4 (Supplementary Table 1). Neoplastic cell clusters were distributed broadly according to MB subgroup, as determined by methylation analysis (Fig. 1a). Neoplastic clusters were most cohesive within the SHH subgroup, with 5 of the 7 SHH samples colocalizing, a reflection of underlying transcriptomic similarity. The 2 discrete SHH clusters were each patient-specific, notably from the two SHH patients with germline *TP53* mutations and particularly extensive CNV aberrations (Supplementary Table 1). GP3 demonstrated the least cohesive clustering, with all patient samples clustering separately. In GP4 5 of 11 samples colocalized. We mapped the sum of CNVs in each sample onto the UMAP which showed a that lower CNV counts were seen in those samples that colocalized within their respective subgroup (Fig. 1c). This finding suggests that sample specific clustering within MB subgroups is a function of the extent of CNV aberration (Fig. 1c). These findings underscore the significant contribution of CNVs to the transcriptional landscape in MB.

**Fig. 1.**
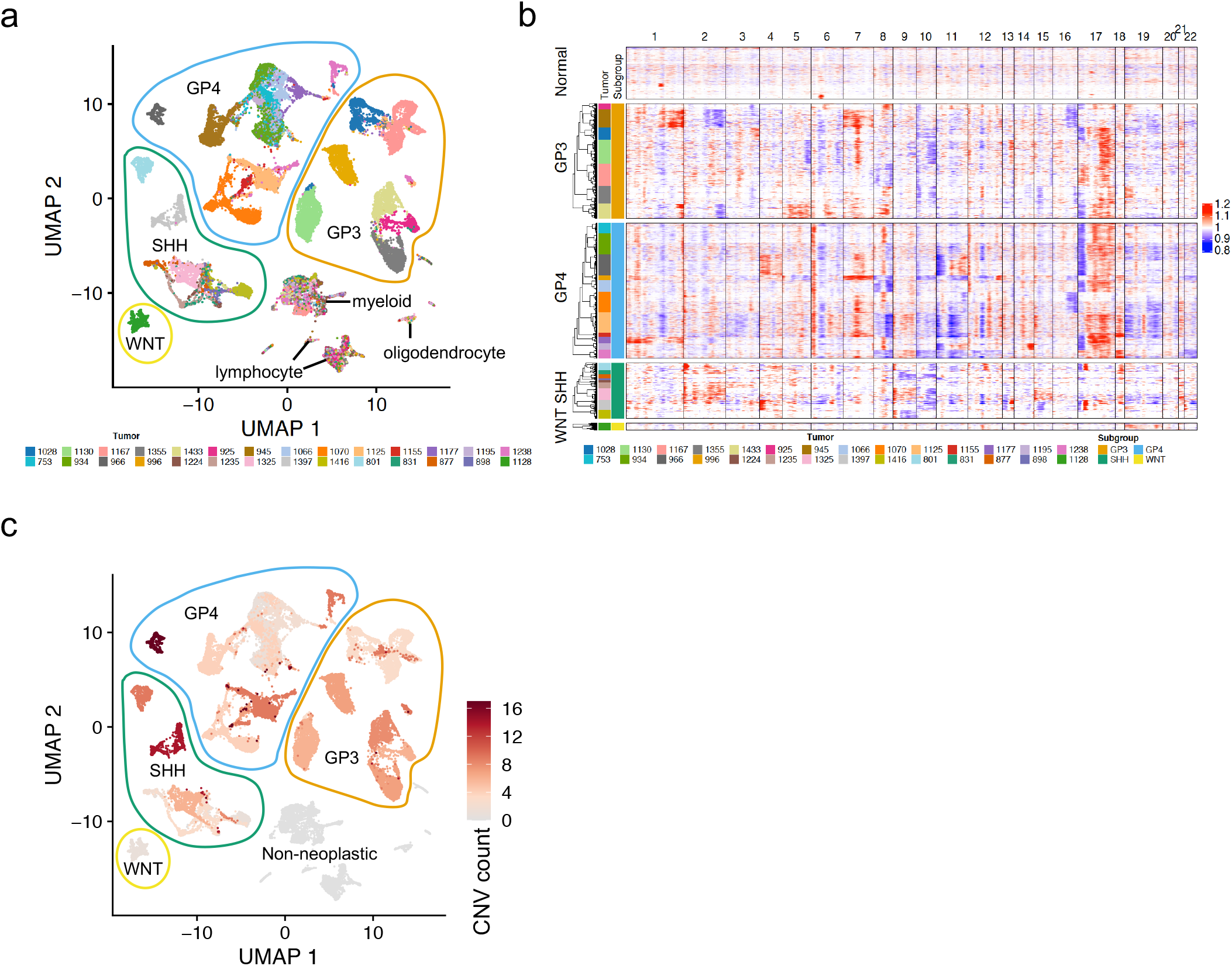
Sample specific clustering in MB scRNA-seq is a function of the extent of chromosomal copy number gain and loss. **a** Unaligned UMAP projection of single-cell expression data of 28 MB patient samples reveals neoplastic clusters and non-neoplastic lymphocyte and myeloid clusters. **b** Inference of CNVs in scRNA-seq data. **c** Copy number gain or loss event count overlaid onto unaligned UMAP projection of MB cells. Abbreviations: astro, astrocyte; oligo, oligodendrocyte; vasc, vascular endothelium.

### GP3 MB contains a differentiated subpopulation corresponding to photoreceptor differentiation

Given the observed subgroup-specific clustering, SHH, GP3 and GP4 subgroup neoplastic cells were individually re-clustered. We used Harmony alignment algorithm to identify transcriptionally similar cell types that had clustered discretely in the prior non-aligned analysis (8). Harmony revealed neoplastic clusters shared between multiple samples in each of the subgroups (Fig. 2–4). SHH, GP3 and GP4 samples each formed clusters that exhibited transcript signatures matching those clusters identified in a recent MB single cell RNA-seq study by Hovestadt *et al.* namely cell cycle (program A), undifferentiated progenitors (program B) and neuronally-differentiated (program C)(5)(Supplementary Fig. 2). We annotated clusters accordingly, but with new subdivisions within these programs corresponding to distinct subpopulations that were identified in the present study. Subpopulations were then characterized by systematic analysis of gene expression, gene ontology (GOTerm) enrichment and inference of active transcription factors (TFs) using pySCENIC (single-cell regulatory network inference and clustering)(9)(Supplementary Tables 3-5, summarized in Table 1). To address similarities of neoplastic subpopulations between MB subgroups, we also performed a combined comparative analysis (Supplementary Fig. 3a).

**Fig. 2.**
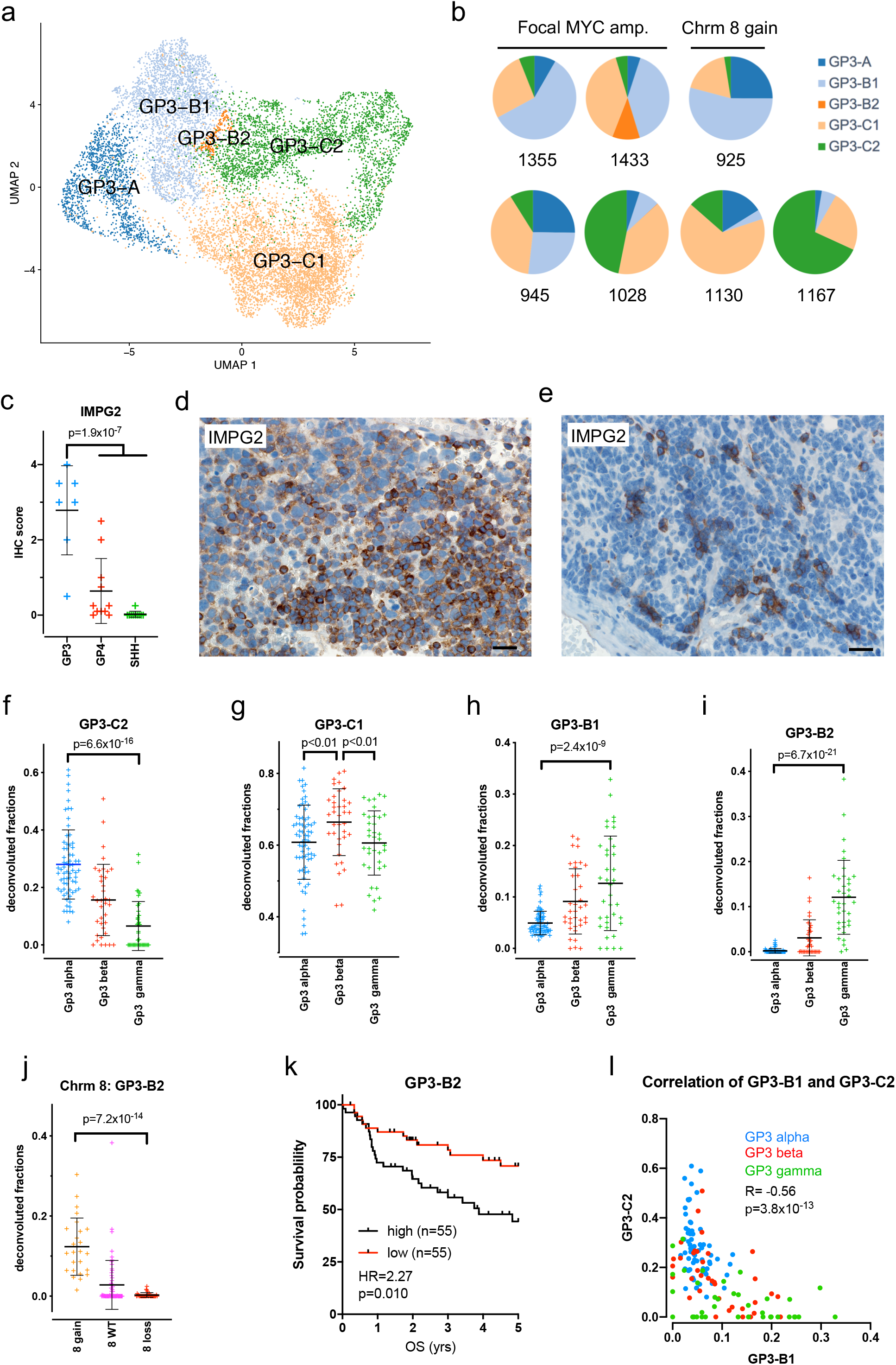
GP3 MB are comprised of progenitor and differentiated neoplastic subpopulations including a photoreceptor-differentiated cluster. **a** Harmony alignment of GP3 neoplastic cells colored by identified clusters. **b** Subpopulation proportions in each GP3 sample. **c** GP3-C2 photoreceptor differentiated subpopulation marker IMPG2 IHC score in MB (GP3, n=7; GP4, n=11; SHH, n=10). Representative IHC staining pattern of IMPG2 (brown) in **d** GP3 (IHC score = 3) and **e** GP4 (IHC score = 2) patient samples (scale bars = 20μm). Deconvoluted **f** GP3-C2 photoreceptor, **g** GP3-C1 neuronally-differentiated, **h** GP3-B2 and **i** GP3-B1 progenitor subpopulation fractions in the MAGIC GP3 cohort (GP3-alpha, n=67; GP3-beta, n=37; GP3-gamma, n=37). Association of **j** chromosome 8 gain, and **k** patient survival with proportion of GP3-B2 (MAGIC GP3 cohort). **l** Subpopulation fraction correlation (Pearson) between GP3-C2 and GP3-B1 (MAGIC GP3 cohort) colored by subtype.

**Table 1.** Summary of MB subgroup neoplastic subpopulation characteristics.

| program |  | GP3 | GP4 | SHH |
| --- | --- | --- | --- | --- |
| **A-cell cycle** |  | **GP3-A** | **GP4-A1** | **SHH-A1** |
|  | GOTerm | cell cycle | cell cycle, G2M, | cell cycle, G2M |
|  | inferred TF activity | MYBL2 | FOXM1, MYBL2 | MYBL2 |
|  | marker genes | CENPF, TOP2A, TUBA1B, HMGB2, MKI67 | TOP2A, HMGB2, CENPF, TUBA1B, MKI67 | CENPF, TOP2A, HMGB2, MKI67 |
|  | proportion | 12.5% | 7.5% | 12.1% |
|  |  |  | **GP4-A2** | **SHH-A2** |
|  | GOTerm |  | cell cycle, S phase | cell cycle: S phase |
|  | inferred TF activity |  | ZNF492, E2Fs | E2Fs |
|  | marker genes |  | HELLS, XRCC2, ATAD5, BRCA1 | RRM2, POLQ, FANCA, XRCC2, BRCA1, HELLS |
|  | proportion |  | 4.6% | 10% |
| **B-progenitor** |  | **GP3-B1** | **GP4-B1** | **SHH-B1** |
|  | GOTerm | translation/oxidative respiration | oxidative phosphorylation/translation | translation/oxidative respiration/neg. reg. of epithelial cell differentiation |
|  | inferred TF activity | HMGA1, MYC, MAX | BARHL1, NEUROD2, SOX4, JUNB, JUND | ATOH1, ZEB1, HES6 |
|  | marker genes | ribosomal | STMN2, ribosomal | SFRP1, CXCR2, ribosomal |
|  | proportion | 33.3% | 25.9% | 28.7% |
|  |  | **GP3-B2** | **GP4-B2** | **SHH-B2** |
|  | GOTerm | translation/oxidative respiration | translation | translation/oxidative respiration |
|  | inferred TF activity | HLX, ATF4, MYC | HEY1, TEAD2, ELK1 | ELK1, THAP11, MAX |
|  | marker genes | ribosomal, MYC | STMN2, ribosomal | ribosomal |
|  | proportion | 1.9% | 17.3% | 17.1% |
| **C-neuronally differentiated** |  | **GP3-C1** | **GP4-C1** | **SHH-C1** |
|  | GOTerm | RNA processing/ axo-dendritic transport | RNA processing/ axo-dendritic transport | RNA processing/ axo-dendritic transport |
|  | inferred TF activity | ARNT | ONECUT2, LHX4 | POU2F2, KMT2A |
|  | marker genes | LUC7L3, DST | GRIA2, LUC7L3, DST | GRIA2, LUC7L3, DST |
|  | proportion | 37.6% | 32.7% | 28.2% |
|  |  | **GP3-C2** | **GP4-C2** | **SHH-C2** |
|  | GOTerm | photoreceptor | photoreceptor | neuron development |
|  | inferred TF activity | SHOX2, CRX | MEIS1, CRX | SOX4, NEUROD1 |
|  | marker genes | NRL, IMPG2 | NRL, IMPG2 | STMN4, STMN2 |
|  | proportion | 14.6% | 6.9% | 3.6% |

Abbreviations: TF, transcription factor.

GP3 tumors (n=7) formed 5 major clusters that included two differentiated cell subpopulations (GP3-C2, -C1) in addition to two progenitor (GP3-B1, -B2) and a mitotic (GP3-A) subpopulation (Fig. 2a,b). Identification of neuronally-differentiated subpopulations in GP3 differs from the findings of Hovestadt *et al.* who reported that GP3 tumors with *MYC* amplification lacked differentiated cells altogether (5). The predominant GP3 differentiated subpopulation GP3-C1 is characterized by genes related to RNA processing and axo-dendritic transport (Table 1, Supplementary Tables 3,4). The less abundant GP3-C2 was distinguished by genes and TFs related to photoreceptor development (*NRL*, *IMPG2*) (Table 1, Supplementary Tables 3-5), a phenotype that has previously been identified and studied in GP3 (10–13). We validated the presence of a photoreceptor-like subpopulation in GP3 using immunohistochemistry (IHC) for interphotoreceptor matrix proteoglycan 2 (IMPG2), a marker of GP3-C2. In a panel of GP3, GP4 and SHH MB formalin-fixed paraffin-embedded (FFPE) samples, IMPG2 protein expression was shown to be significantly higher in GP3 than GP4 and SHH (p=1.9×10^−7^), with an expression score of 2 or greater in 6/7 GP3 samples (Fig. 2c). IMPG2 was expressed in discrete clusters of cells in GP3 tumors, suggesting focal areas of GP3-C2 photoreceptor differentiation (Fig. 2d). Discrete clusters of IMPG2 expressing cells were also observed in a smaller number of GP4 tumors (Fig. 2e). The patchy distribution of *IMPG2*, and presumably other photoreceptor subpopulation-restricted markers, across areas of tumor in both GP3 and to a lesser extent in GP4 could therefore introduce a confounding factor for classification techniques that employ these markers for GP3 (14)(Supplementary Fig. 4).

Subpopulation proportions were estimated by deconvolution (Cibersort) of the Medulloblastoma Advanced Genomics International Consortium (MAGIC) dataset, a large patient cohort that includes transcriptomic data, subgroup/subtype classification and associated CNV and outcome data for the majority of samples(n=747) (2). For individual MB subgroup, estimated subpopulation proportions were compared between samples grouped according to subtype, CNVs and clinical outcome to identify significant associations. Deconvolution analysis of the GP3 MAGIC cohort (n=141) revealed a significant association between photoreceptor-differentiated subpopulation GP3-C2 and subtype GP3 alpha, that has the best clinical outcome of all GP3 subtypes per Cavalli *et al.* (2). The GP3-C2 proportion was 2.5-fold higher in GP3 alpha than other GP3 subtypes (p=2.3×10^−14^), being progressively less abundant in GP3 beta and GP3 gamma (Fig. 2f). GP3 differentiated subpopulation GP3-C1 was more abundant in GP3 beta than others (p=0.0027)(Fig. 2g).

### GP3 progenitor proportions are influenced by MYC and chromosome 8 gain

Examination of GP3 progenitor subpopulations GP3-B1 and GP3-B2 further substantiated the previously articulated theory that aberrant *MYC* activity is responsible for preventing differentiation of GP3 progenitor subpopulations (5). Both GP3-B1 and B2 shared GOTerm enrichment of translation-related genes, including ribosomal proteins and eukaryotic translation elongation factors, and MYC TF activity (Table 1, Supplementary Tables 3-5). One normal physiological function of MYC is to directly enhance Pol I transcription of ribosomal RNA (rRNA) genes in response to the cellular demand for protein synthesis (15, 16). This normal physiological role of MYC is recapitulated in the GP3-B progenitor cells, and focal amplification of *MYC* or gain of chromosome 8 is associated with increased proportions of progenitor cells versus non-*MYC* amplified GP3 samples (p=0.013)(Fig. 2b). Deconvolution of the MAGIC GP3 cohort showed a significantly higher proportion of progenitor GP3-B1 and GP3-B2 cells in GP3 gamma than in either alpha or beta subtypes (Fig. 2h,i). GP3-gamma samples are distinguished by broad copy number gains on chromosome 8, focal copy number gains for *MYC* and the poorest clinical outcome of all MB subtypes (2). We compared all subpopulation proportions to broad CNVs reported for the MAGIC sample cohort, which showed that the strongest of these associations was between GP3-B2 and chromosome 8 CNVs, with GP3-B2 proportions being higher in samples with chromosome 8 gain than in samples with chromosome 8 WT or loss (Fig. 2j). These data are consistent with the higher proportions of GP3 progenitor subpopulations in our scRNA-seq samples with gain of chromosome 8 or *MYC* amplification (Fig. 2b). With respect to clinical outcome, a higher than median proportion of GP3-B2 conferred the shortest survival of all GP3 subpopulations (Fig. 2k, Supplementary Table 6), consistent with the poor outcome conveyed by Gp3-gamma subtype assignment and *MYC* amplification.

The relationship between deconvoluted GP3 subpopulation proportions was examined, identifying a significant negative correlation between GP3-C2 and all other GP3 subpopulations in particular GP3-B1 (Fig. 2l, Supplementary Table 7). This analysis demonstrates a continuum of differentiated and progenitor cell ratios across GP3 alpha, beta and gamma subtypes, an important factor for correct interpretation of GP3 biological studies.

### Chromosome 8 loss is associated with lower proportion of GP4 progenitors

Six major neoplastic subpopulations were identified in 11 GP4 scRNA-seq samples, with some degree of similarity with subpopulations identified in GP3 (Fig. 3a,b; Table 1). Two subpopulations were characterized by cell cycle-related gene enrichment (GP4-A1 and A2). We identified 2 progenitor subpopulations (GP4-B1 and -B2) that were characterized by abundant ribosomal gene expression, as seen in GP3 progenitors (Table 1, Supplementary Tables 3,4). Deconvolution analysis of GP4 MAGIC samples (n=325) revealed that the strongest association between CNVs and estimated subpopulation proportions was between GP4-B1 and chromosome 8 aberrations (Fig. 3c). This association differs from that seen with in GP3 in that GP4-B1 proportions are significantly higher in samples with WT chromosome 8 than those with chromosome 8 loss (chromosome 8 gain being largely absent in GP4 MB). This finding implicates a role for MYC in the maintenance of a progenitor subpopulation in GP4 as well as GP3 and warrants further investigation. GP4 subtypes also showed variable subpopulation proportions, the most distinct being GP4 gamma having significantly lower GP4-B1 progenitors (p=3.2×10^−18^)(Fig. 3c).

**Fig. 3.**
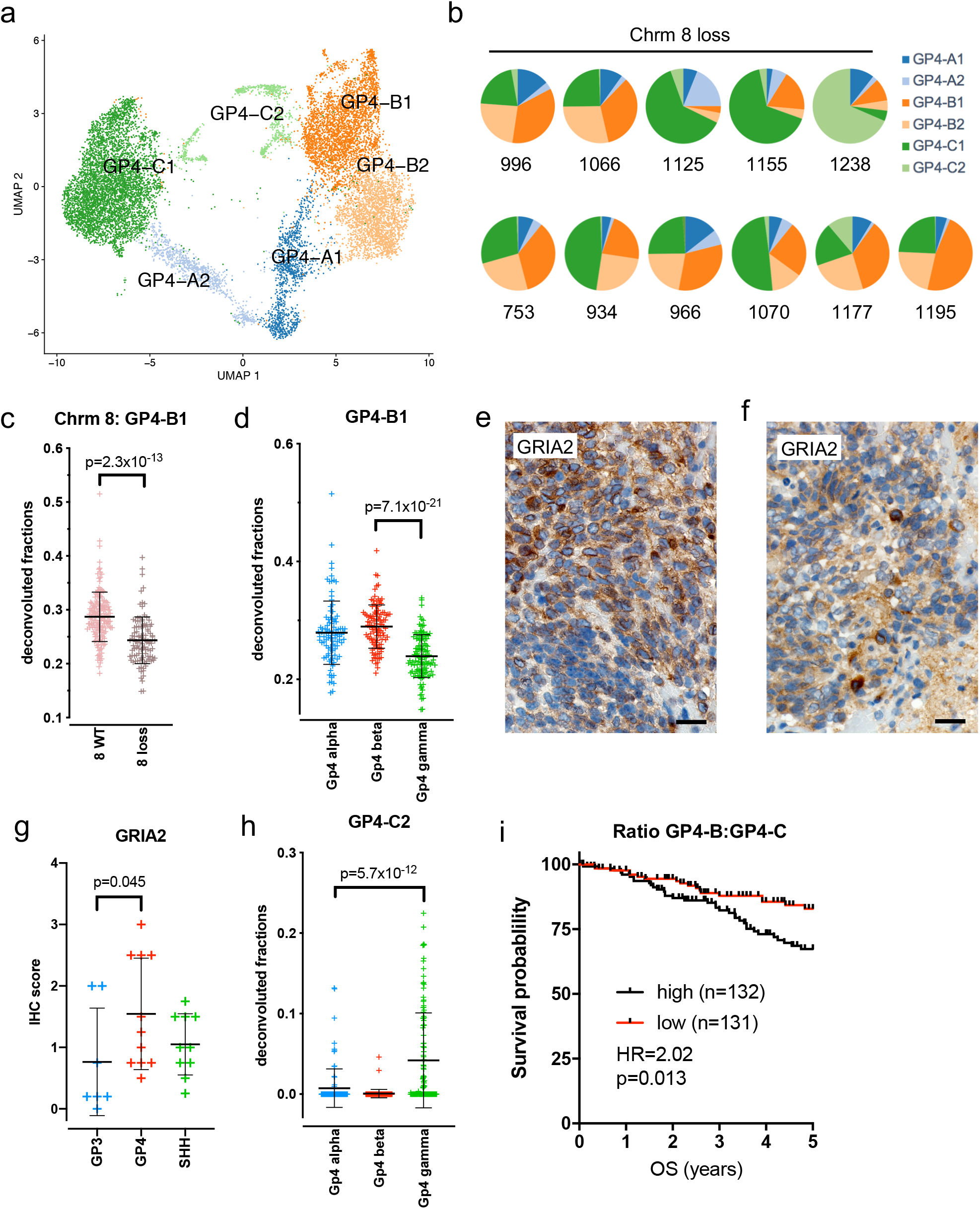
GP4 MB neoplastic cell heterogeneity. **a** Harmony alignment of GP4 neoplastic cells colored by identified clusters. **b** Subpopulation proportions in each GP4 sample. **c** Association of chromosome 8 loss, and **d** subtype with deconvoluted proportion of GP4-B1 (MAGIC GP4 cohort: GP4-alpha, n=97; GP4-beta, n=109; GP4-gamma, n=11). Representative IHC staining pattern of neuron differentiated subpopulation marker GRIA2 (brown) in **e** a GP4 patient sample (IHC score = 2.5) and **f** a GP3 patient sample (IHC score = 0.75)(scale bars = 20μm). **g** GP4-C2 subpopulation marker GRIA2 IHC score in a cohort of MB (GP3, n=7; GP4, n=11; SHH, n=10). **h** Deconvoluted GP4-C2 photoreceptor subpopulation fractions (MAGIC GP4 cohort). **i** Association of patient survival with ratio of progenitor to differentiated deconvoluted fractions (MAGIC GP4 cohort).

### The predominant GP4 MB differentiated subpopulation is distinguished by expression of glutamatergic neuron lineage markers

The predominant differentiated GP4 subpopulation, GP4-C1, is characterized by gene expression similar to GP3-C1, including expression of glutamate receptor *GRIA2*, a key component of glutamatergic lineage neurons (Supplementary Table 3). In recent single cell analyses of the developing human cerebellum *GRIA2* was a marker of glutamatergic lineage neurons, most notably human excitatory cerebellar nuclei (eCN)/uniopolar brush cells (UBC) (17), although also expressed at lower levels in Purkinje cells and cerebellar granular neurons (CGNs)(18). We compared human cerebellar lineage scRNA-seq data (17) with our MB neoplastic subpopulations showing that overlap between these cerebellar cell types was largely restricted to MB differentiated subpopulations and absent from progenitor subpopulations (Supplementary Fig. 5a-c). The strongest correlation was observed between human eCN/UBC signatures and GP4-C1, and to a progressively lesser extent with SHH-C1 and GP3-C1 (Supplementary Fig. 5d). Accordingly, IHC analysis revealed that eCN/UBC marker GRIA2 positive cells were abundantly distributed throughout the tumor parenchyma in GP4 (Fig. 3e), to a moderate extent in SHH and least abundant in GP3 (Fig. 3f,g). These findings are consistent with early MB bulk-tumor transcriptomic data that revealed a glutamatergic signature in GP4 and also in GP3, but to a lesser extent (10). The same study identified a GABAergic signature that was exclusive to GP3. Although no MB neoplastic subpopulations overlapped with GABAergic lineages, GABRA5, a key marker of the GP3 GABAergic signature (10) was expressed predominantly in GP3 progenitors and GP3-C1 (Supplementary Fig. 4a).

A photoreceptor differentiated subpopulation, GP4-C2, was also variably observed in GP4, although to a lesser extent than in GP3 (Fig. 3b, Table 1). Comparison of GP4-C2 with other neoplastic subpopulations confirmed the strong correlation with GP3-C3 (Supplementary Fig. 3a). Photoreceptor-differentiated cells in GP4 was examined by IHC (Fig. 2e), with 2/11 GP4 samples expressing IMPG2 with an IHC score of 2 or more, and isolated IMPG2 positive cells in the remainder (Fig. 2c). The presence of photoreceptor-differentiated cells in GP4 may explain those MB samples that are assigned to the mixed GP3/GP4 category, as illustrated by the expression of GP3 nanostring markers IMPG2 and NRL (14) in the GP4-C2 subpopulation (Supplementary Fig. 4a). By deconvolution analysis, MAGIC cohort GP4 subtypes alpha, beta and gamma also showed variable differentiated subpopulation proportions, with a significantly higher photoreceptor differentiated subpopulation GP4-C2 in GP4 gamma (11-fold higher than others, p=5.7×10^−12^)(Fig. 3h).

No single GP4 subpopulation was significantly correlated with survival (Supplementary Table 6), although the ratio of progenitor (GP4-B1 and -B2 combined) to differentiated (GP4-C1 and -C2 combined) conveyed a shorter survival (Fig. 3i). These findings suggest that the better clinical survival reported for GP4-gamma compared to other GP4 subtypes (2) is likely attributable to a higher ratio of differentiated to progenitor cells.

### SHH tumors contain a unique differentiated subpopulation that comprises nodular regions

We identified six major subpopulations of neoplastic cells in SHH tumors (n=9) including 2 cell cycle (SHH-A1 and -A2), 2 progenitor (SHH-B1 and -B2) and 2 neuronally-differentiated subpopulations (SHH-C1 and -C2) (Fig. 4a,b). SHH-C2 was distinct from SHH-C1 and differentiated subpopulations in other subgroups (Supplementary Fig. 3a) with a particularly high enrichment of neuron-related gene ontologies (Fig. 4c, Supplementary Table 4). SHH-C2 was distinguished from SHH-C1 by high expression of stathmin genes (*STMN2* and *STMN4*) which are involved with neuron development and growth (19). IHC showed that SHH-C2 marker STMN2 positive cells were predominant in SHH nodules (Fig. 4d,e). Nodules of neuronal maturation are a classic histologic feature of SHH that recapitulates aspects of normal cerebellar development (20). In the MAGIC cohort, Cavalli *et al.* delineated four SHH subtypes: two infant subtypes, SHH beta and SHH gamma (the latter largely comprised of MB with extensive nodularity (MBEN)); a subtype in older children, SHH alpha which harbors frequent p53 mutations; and a largely adult subtype, SHH delta (2). We deconvoluted the MAGIC SHH transcriptomic cohort (n=214) and showed that SHH-C2 was the most differential subtype distribution being significantly more abundant in the infant SHH beta and gamma subtypes (Fig. 4f). Consistent with SHH-C2 representing the primary population of cells that constitute SHH nodules, this subpopulation was most abundant in highly nodular SHH gamma that is associated with SHH variant MBEN.

**Fig. 4.**
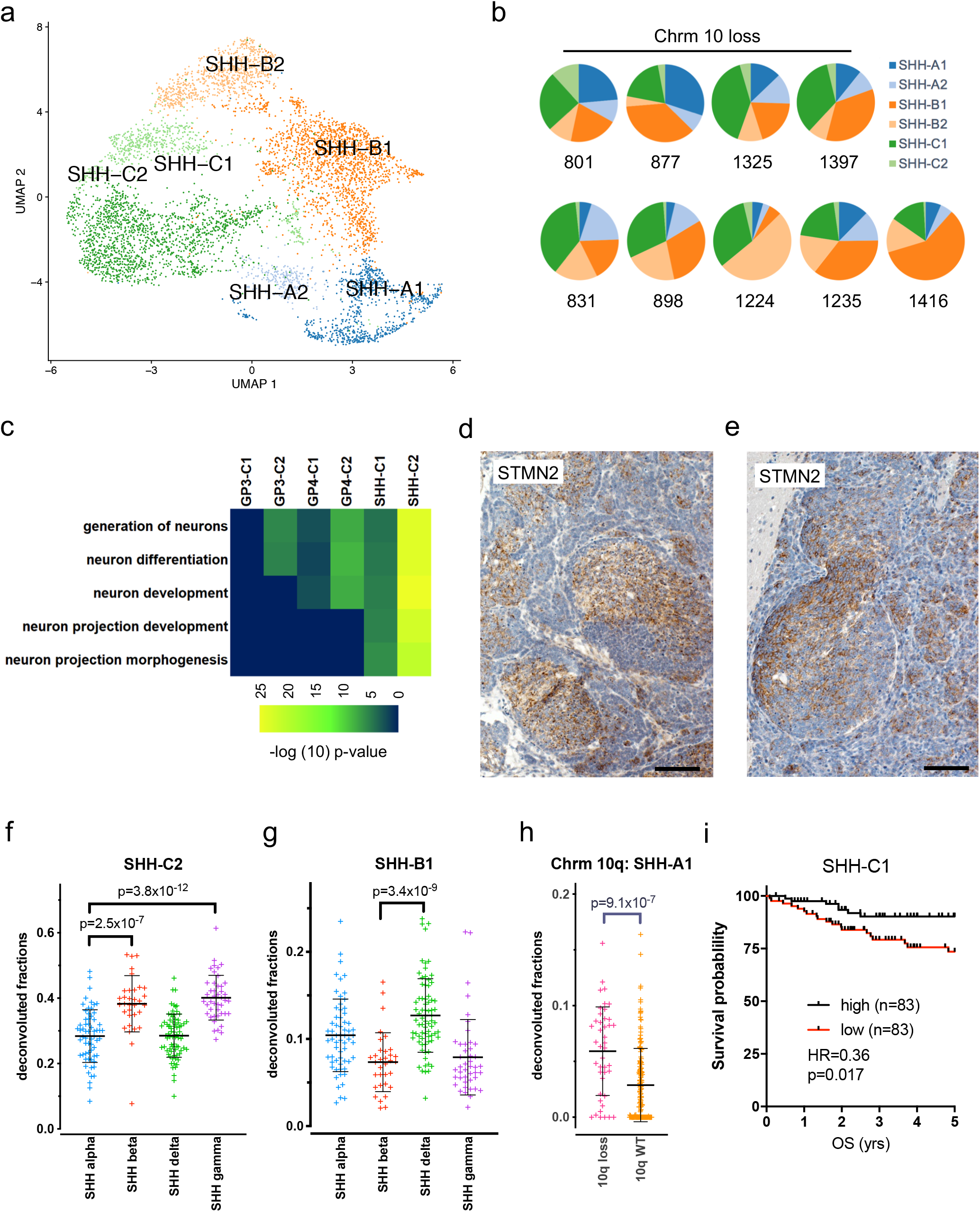
SHH MB neoplastic subpopulations include a nodule-associated neuronally-differentiated subpopulation. **a** Harmony alignment of SHH neoplastic cells colored by identified clusters. **b** Subpopulation proportions in each SHH sample. **c** Heatmap showing enrichment of top neuron-related GOTerms across differentiated subpopulations. **d,e** Representative IHC staining patterns of SHH-C2 subpopulation marker STMN2 (brown) in a SHH patient samples (scale bars = **d** 40μm and **e** 20μm). Deconvoluted **f** SHH-C2 nodule-associated, and **g** SHH-B1 progenitor subpopulation fractions (MAGIC SHH cohort: SHH-alpha, n=62; SHH-beta, n=33; SHH-delta, n=75; SHH-gamma, n=44). **h** Association of chromosome 10q loss with deconvoluted proportion of GP4-A1 (MAGIC SHH cohort). **i** Association of patient survival with deconvoluted SHH-C1 neuron-differentiated fraction (MAGIC SHH cohort).

The most abundant neuronally-differentiated SHH subpopulation SHH-C1 was most closely correlated with GP4-C1 and GP3-C1 (Supplementary Fig. 3a), with enrichment of RNA processing and axo-dendritic transport ontologies (Table 1, Supplementary Tables 3,4) and overlap with human glutamatergic eCN/UBC lineage gene expression (Supplementary Fig. 5). SHH-C1 was the only SHH subpopulation significantly associated with outcome, its relative abundance being favorable for survival (Fig. 4i, Supplementary Table 6).

SHH progenitors are distinct from GP3 and GP4 by expression of previously identified components of SHH biology. These include inferred TF activity of CGN progenitor specification gene ATOH1 (Supplementary Table 5) and high expression of genes associated with negative regulation of epithelial cell proliferation and WNT signaling (*SFRP1*, *SFRP5*) (Supplementary Tables 3,4). This is consistent with studies showing that WNT pathway activation is antagonistic to SHH-driven proliferation (21, 22). SHH progenitor subpopulations SHH-B1 and -B2 were similar to GP3 and GP4 progenitor subpopulations with respect to abundant ribosome and eukaryotic translation elongation factor gene expression (Supplementary Tables 3,4). The *MYCN* gene is commonly amplified in SHH, and like cMYC can upregulate ribosomal RNA expression as a mechanism of universal upregulation of gene expression (23). Related to this, germline elongation factor *ELP1* mutations have recently been implicated as a major driver in SHH etiology (24), further supporting a potential role of transcriptional/translational dysregulation in SHH progenitor biology. SHH progenitor populations, in particular SHH-B1, were significantly inversely correlated with SHH-C2 (Supplementary Table 7), being highest in non-infant subtypes SHH alpha and delta (Fig. 4g).

CNVs were not as strongly correlated with SHH subpopulations as in GP3 or GP4, the most significant being an increase (2.1-fold higher) in cell cycle subpopulation SHH-A1 in those samples with 10q loss (Fig. 4h). This supports a role of PTEN loss in upregulation of cell cycle in SHH that has was also observed in a *PTEN*-deficient SHH mouse model (25). SHH-A subpopulations, and GP3-A and GP4-A, were enriched for cells in both G2M and S-phase, although we note that other subpopulations contain proliferating cells to a lesser extent (Supplementary Fig. 3b).

### The immune cell compartment of MB contains diverse myeloid cell subpopulations associated with neurodevelopment

ScRNA-seq provided us with an opportunity to comprehensively chart tumor infiltrating immune cell subpopulations in MB. Cells identified as lymphocyte or myeloid (n=4669), were separated out from neoplastic cells and re-clustered, revealing four lymphocyte, six myeloid cell and a single cell cycle-related cluster (Fig. 5a). Myeloid and immune cells from multiple patients’ samples strongly co-clustered and therefore did not require Harmony alignment.

**Fig. 5.**
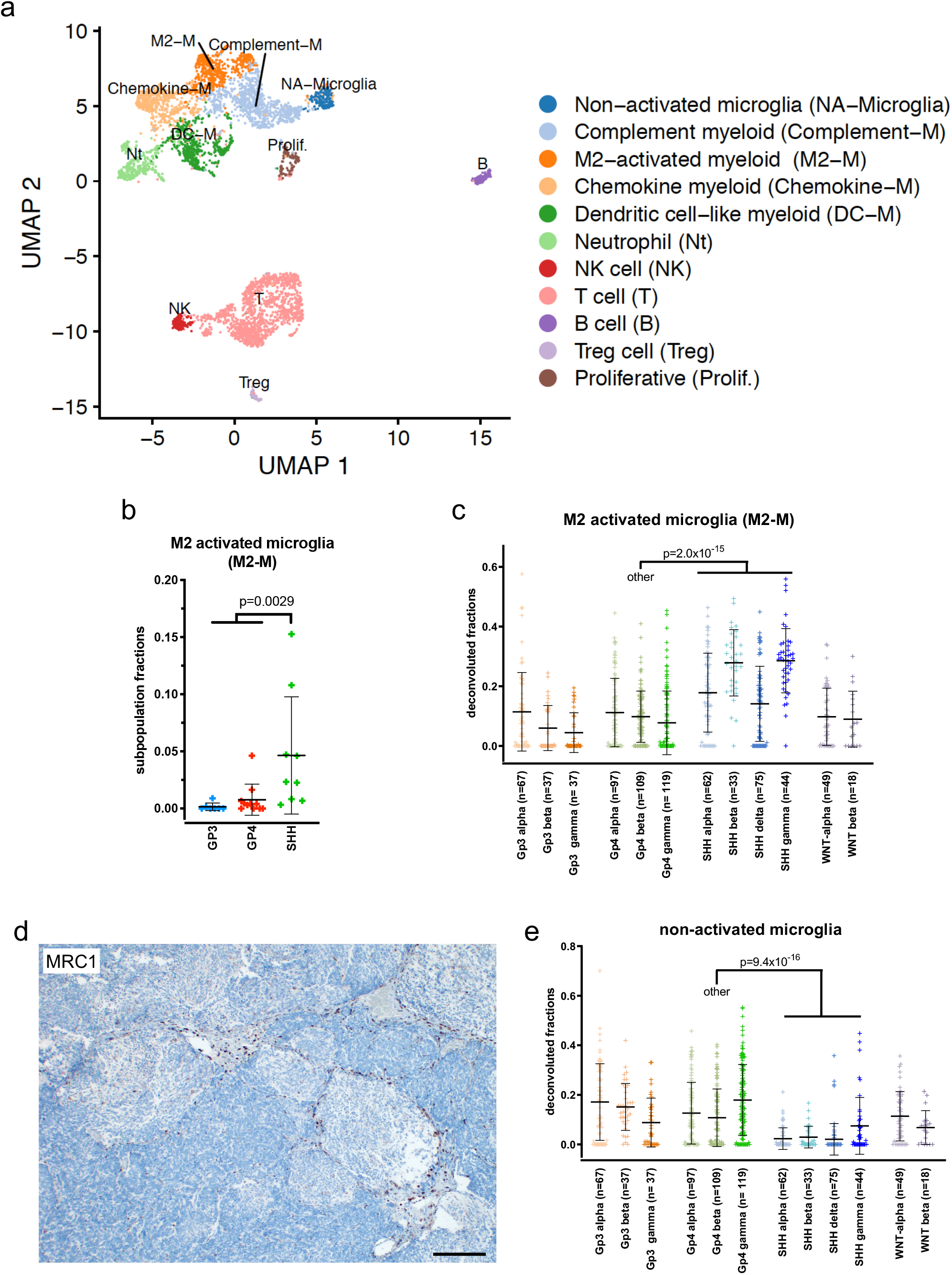
The immune landscape of MB. **a** non-harmonized alignment of MB tumor infiltrating immune cells colored by identified clusters. **b** M2-myeloid subpopulation fractions (scRNA-seq cohort; GP3, n=7; GP4, n=11; SHH, n=9). **c** Deconvoluted M2-myeloid (M2-M) subpopulation in MB subtypes (MAGIC cohort). **d** Representative IHC staining pattern of M2-myeloid subpopulation marker MRC1 (brown) in an SHH patient sample showing accumulation at nodule linings (scale bar = 100μm). **e** Deconvoluted non-activated microglia subpopulation in all MB subtypes (MAGIC cohort).

An unexpected degree of myeloid cell diversity was observed in the majority of MB samples analyzed (Fig. 5a). This finding is consistent with other scRNA-seq analyses of tumor infiltrating and CNS immune subpopulations that showed a greater degree of diversity than was previously assumed (26, 27). Despite apparent transcriptional overlap between subpopulations identified in these recent studies no consensus nomenclature yet exists. Accordingly, immune subpopulations described in the present populations are annotated with names descriptive of transcriptional signatures. Myeloid populations in the CNS are separated into two main lineages - tissue-resident microglia and peripheral bone marrow-derived myeloid cells – and both of these play roles in both immune response and neurodevelopment (28). These shared functional roles have hindered the identification of distinguishing markers for the two etiological lineages. Consequently, we describe MB myeloid subpopulations according to neurodevelopmental or immune processes inferred from transcriptomic signatures (Supplementary Table 8) and cannot for the most part reliably assign subpopulations to resident or peripheral myeloid origins.

The most abundant myeloid subpopulation exhibited signatures associated with neurodevelopmental roles. Specifically this subpopulation demonstrated particularly high expression of all complement component 1q subunits (*C1QA*, *C1QB* and *C1QC*) (Supplementary Table 8), that are responsible for the initiation of the classic complement cascade that is involved in neurodevelopmental synapse elimination (29). This population was thus termed complement myeloid (Complement-M). A second myeloid neurodevelopment-associated subpopulation was characterized by expression of phagocytosis receptors *MRC1* (CD206), CD163, and *MERTK* that are crucial for microglia-mediated clearance of apoptotic cell debris from the developing neurogenic niche (30). *MRC1* and *CD163* are widely recognized markers of anti-inflammatory or immunoregulatory M2 myeloid polarization and this population was therefore named M2-activated myeloid (M2-M). M2-M cells showed the strongest subgroup association, being more abundant (8.8-fold) in SHH than GP3 and GP4 combined in scRNA-seq samples (Fig. 5b). Myeloid cell scRNA-seq subpopulation signatures were used to deconvolute the MAGIC dataset (GP3, GP4, SHH and WNT; n=748) which corroborated the observation that M2-M were more abundant in SHH that other subgroups (2.2-fold; p=2.0×10^−15^) (Fig. 5c). Further, M2-M abundance was significantly higher in infant subtypes SHH beta and gamma than older subtypes SHH alpha and delta (1.8-fold; p=3.7×10^−6^)(Fig. 5c). IHC for M2-M marker *MRC1* (CD206), stained cells with an amoeboid morphology that frequently lined SHH nodules (Fig. 5d). The co-localization of M2-M with nodules, which are comprised of the strongly neuronally-differentiated SHH-C2 subpopulation cells support the hypothesis that this myeloid subpopulation is programmed for developmental activity, such as synaptic pruning and elimination of apoptotic neuron debris, rather than an immune role. This hypothesis is also supported by age restriction of M2-M, being significantly more abundant in infant SHH subtypes that are likely more developmentally influenced.

A single myeloid subpopulation could confidently be designated as microglia, expressing widely established microglial markers (*P2RY12, TMEM119, SALL1, CX3CR1*)(Supplementary Table 8). Due to the relatively low expression of myeloid cell immune activation markers (including HLA class II and cytokines/chemokines) this myeloid population was classified as non-activated microglia (NA-microglia). In contrast to M2-M, NA-microglia trended toward being less abundant in SHH compared to GP3 and GP4 in scRNAseq samples and were significantly less abundant in SHH in the larger MAGIC cohort (4-fold, p=9.4×10^−16^)(Fig. 5e). These findings are congruent with prior studies that identified a less activated immune phenotype in GP3 and GP4 than SHH (31, 32).

Two further myeloid subpopulations exhibited transcriptomic profiles associated with antigen presentation, both expressing MHC class II genes, suggesting an active immune functional role as opposed to the developmental/homeostatic roles inferred for the prior myeloid subpopulations. The first of these was distinguished from previous myeloid subpopulations by expression of chemokines (*CCL3, CCL4, CXCL8* and *IL1B*) and MHC class II (*HLA-DRB1*, *HLA-DRA, HLA-DPA*)(Supplementary Table 8), suggesting immune cell recruitment and antigen presentation functions, respectively, and is referred to as chemokine myeloid (Chemokine-M). A second antigen presenting myeloid subpopulation was distinguished by C-lectins (*CLEC10A, CLEC4A, CLEC12A*), *CD1* subunits (*CD1C, CD1D* and *CD1E*) and MHC class II (*HLA-DQA1, HLA-DQB1*) all of which are associated with antigen presentation by macrophages and/or dendritic cells, and was therefore named dendritic cell-like myeloid (DC-M). Outcome analysis of deconvoluted myeloid immune subpopulation factions did not identify any significant association with survival for antigen presentation-associated myeloid subpopulations Chemokine-M or DC-M (Supplementary Table 6). A survival advantage was instead observed in those GP3 and GP4 with a higher than median proportion of complement-M cells, suggesting a significant role for myeloid cells associated with putative neurodevelopmental processes in MB biology.

Analysis of lymphocytes in MB was limited by their rarity in this tumor type compared to other pediatric brain tumors, including glioblastoma, as has previously been observed (33). The most abundant MB lymphocyte population was identified as T-cells (*CD3D, TRAC*), constituting ~80% of lymphocytes, with no separation of CD4 from CD8 T-cells at this level of resolution (Fig. 5a). The remaining lymphocyte clusters were NK cells (*NKG7, GNLY*), B-cells (*MS4A1*, *CD79A*) and regulatory T-cells (*FOXP3, CTLA4*). None of these lymphocyte subpopulations was significantly differentially distributed across MB subgroups in the scRNA-seq cohort. Deconvolution of the larger MAGIC cohort did not yield usable results, likely due to the scarcity of lymphocyte specific transcripts in bulk transcriptomic data.

### Genetically engineered mouse models of MB contain subpopulations corresponding to human subgroup-specific subpopulations

Genetically engineered mouse models (GEMMs) for GP3 and SHH MB provide valuable *in vivo* experimental models. We sought to define the cellular heterogeneity present in these with respect to identified human MB subpopulations, which is critical for correct interpretation of GEMM experimental results. Two GP3 models, the first (termed MP) driven by overexpression of cMyc and dominant negative *Trp53* (34, 35), and the second driven by co-expression of Myc and Gfi1 (and termed MG)(36, 37), and one SHH model driven by mutant Smo activated in the Atoh1 lineage (Math1-Cre/SmoM2 - termed MS)(38) were examined using scRNA-seq. Harmony was used to align GEMM with human MB subgroup scRNA-seq data, revealing that GEMM cells clustered most closely their intended corresponding human MB subgroups (Fig. 6a,b).

**Fig. 6.**
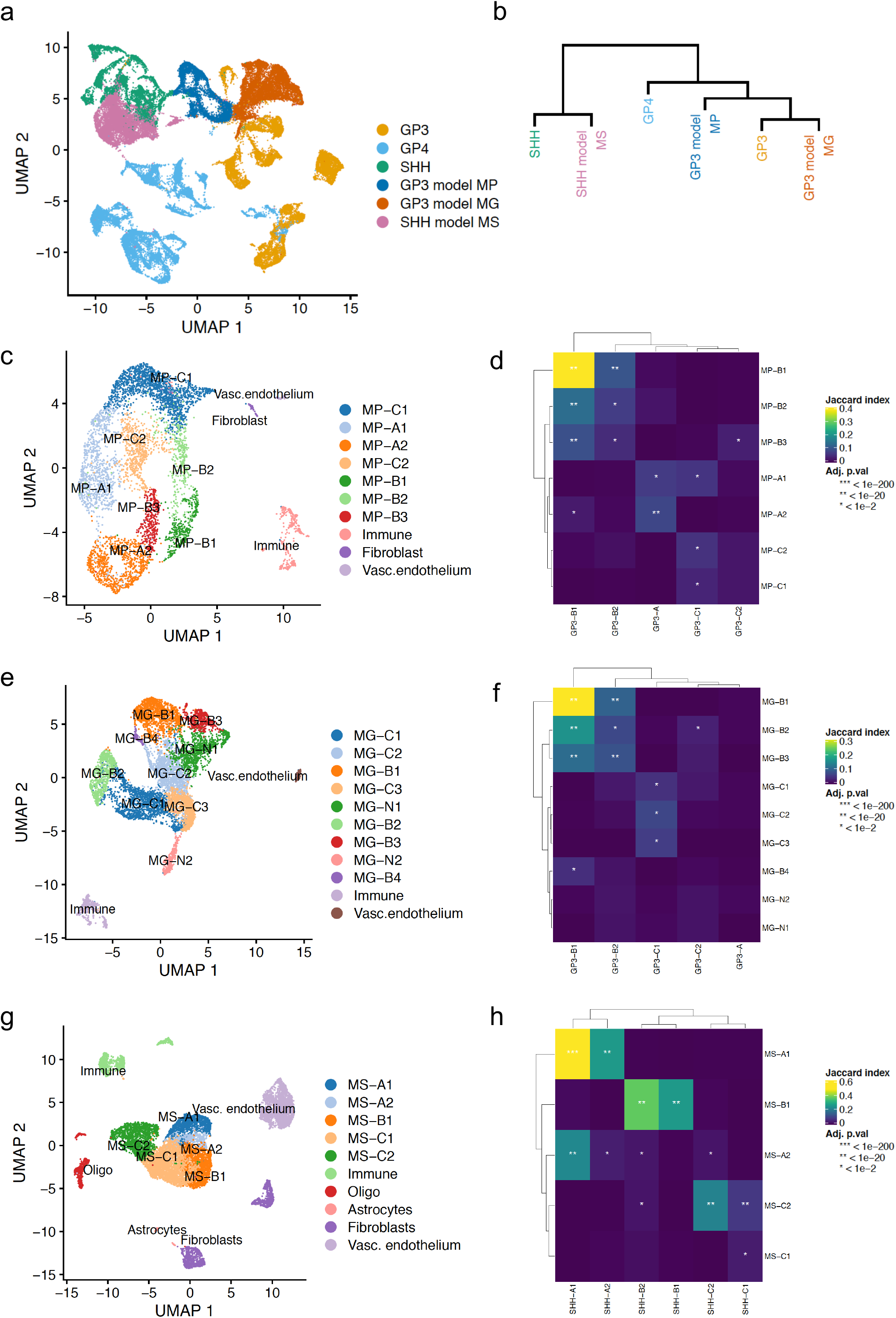
GEMM cellular heterogeneity and fidelity with human MB subpopulations Alignment of GEMM with human MB subgroup scRNA-seq data, revealing that GEMM cells clustered most closely their intended human MB subgroups. **a** UMAP projection of Harmony aligned (theta=2) GEMM and human MB scRNA-seq data. **b** Hierarchical clustering of GEMM and human MB scRNA-seq data. **c** Non-harmonized alignment of GEMM model MP single cells colored by identified clusters, and labelled according to corresponding subgroup human neoplastic subpopulations, and **d** Jaccard index for the top 200 marker genes between GEMM model MP and human GP3 neoplastic subpopulations. This was repeated for **e,f** GP3 GEMM model MG, and **g,h** SHH model MS. Abbr: MP, GP3 GEMM Myc/Trp53; MG, GP3 GEMM Myc/GFI1; MS, SHH GEMM Math-1/SmoM2; vasc., vascular; oligo, oligodendrocyte.

Both GP3 GEMMs were generated by retroviral overexpression of oncogenes in neural progenitors, and transplantation of these progenitors into the cerebellum of adult NSG mice. In model MP, scRNA-seq of 5,422 cells identified 7 neoplastic and 2 non-neoplastic subpopulations (immune and fibroblast) (Fig. 6c). The 7 neoplastic GEMM subpopulations were compared to human GP3 neoplastic subpopulations using three methods: (i) comparison of the top 200 marker genes for GEMM neoplastic subpopulations (Supplementary Table 9) compared to corresponding subgroup subpopulations (Fig. 6d) as used previously to compare neoplastic subpopulations between subgroups; (ii) UMAPs of GEMM cells labeled with human neoplastic subpopulation metagene signatures (Supplementary Fig. 6a); and (iii) correlation of the genome-wide mean expression of each GEMM neoplastic subpopulation with human neoplastic subpopulations which are then visualized using a directed bipartite graph (Supplementary Fig. 6b)(39). Results of these analyses were used to assign GEMM subpopulations to cell cycle (A), progenitor (B) and differentiated (C) phenotypes. This approach identified MP subpopulations corresponding to human GP3 cell cycle (MP-A1, −2), progenitor (MP-B1,-B2, -B3) and differentiated neoplastic subpopulations (MP-C1, -C2) (Fig. 6c). Similarly, the second GP3 GEMM MG harbored neoplastic and non-neoplastic subpopulations (8,144 cells) (Fig. 6e), which were comprised of 9 neoplastic clusters that grossly corresponded to progenitor (MG-B1, -B2, -B3, -B4) and differentiated subpopulations (MB-C1, -C2, -C3) (Fig. 6f, Supplementary Fig. 7, Supplementary Table 9), but with two subpopulations with no apparent human equivalent (MB-N1, -N2) and no apparent cell cycle subpopulations. Both GP3 GEMMs harbored subpopulations corresponding to the neuronally-differentiated human GP3-C1, but with only a very minor subpopulation of cells that were enriched for photoreceptor differentiated GP3-C2 metagenes (Supplementary Fig. 6a, 7a). Collectively, these data suggest that the MP GEMM caries more fidelity to human GP3 tumors SHH GEMM model MS was comprised of subpopulations that also corresponded to human SHH neoplastic subpopulations. Five neoplastic subpopulations and 8 non-neoplastic subpopulations were revealed by scRNA-seq analysis (12,421 cells; Fig. 6g). Neoplastic cells showed strong concordance with human SHH neoplastic subpopulations, comprising 2 cell cycle (MS-A1, −2), 1 progenitor (MS-B1) and 2 differentiated subpopulations (MS-C1, -C2). (Fig. 6h, Supplementary Fig. 8, Supplementary Table 9), including a neuronally-differentiated subpopulation (MS-C2) corresponding to the human SHH nodule-associated SHH-C2 subpopulation. Cumulatively we show that the GEMM MB models are consistent with human MB and can be used to test therapeutic response and resistance to novel agents.

### Human and mouse MB cell atlas browser

We have created a publicly available scRNA-seq resource, the Pediatric Neuro-oncology Cell Atlas (pNeuroOncCellAtlas.org). This interactive resource allows users to study expression of transcripts at the single cell level for neoplastic and immune cells in human samples and GEMM models, with extensive annotations.

## Discussion

Recognition of the inherent intra-tumoral cellular heterogeneity in MB is critical for the correct interpretation of MB cancer biology, an aggressive childhood brain tumor that has seen few treatment advances in 20 years. The present study addresses this problem using scRNA-seq to study childhood MB, revealing neoplastic and immune subpopulations that had not previously been identified. We characterize these novel neoplastic and immune subpopulations to provide deeper insights into a broad range of aspects of MB biology.

Our study identifies associations between chromosomal aberrations both with single cell clustering patterns and progenitor proportions. We show that separation of sample-specific cell clusters is driven in part by the extent of chromosomal copy number gain and loss. Copy number variance of chromosome 8 correlated with the proportion of progenitor populations in GP3, as previously observed, but this association is also observed in GP4. These findings underscore the important role that CNVs play in MB cellular heterogeneity and their potential impact on subgroup/subtype assignment. MB cellular heterogeneity may confound tumor sub-classifications that are based on sequencing of bulk-tumor samples. Several international consortium studies have interogated MB bulk-tumor samples, identifying consensus molecular subgroups with further subdivision of these into multiple subtypes (2–4). Our study reveals differential subpopulation proportions between MB subgroups and subtypes, and restriction of a number of consensus subgroup transcriptional markers to discrete subpopulations. These findings underscore the significant contribution of MB cellular heterogeneity to subgroup/subtype assignment.

Our analysis revealed the presence of a photoreceptor-differentiated subpopulation that was seen predominantly in GP3 and to a lesser extent in GP4 tumor samples. This finding in GP3 samples is concordant with numerous prior studies that had identified a photoreceptor gene signature in MB (12) and more recently as a defining feature of GP3 (10, 11). The proportion of photoreceptor differentiated cells was variable in GP3, being inversely proportional to MYC-associated progenitor subpopulations. In the MAGIC transcriptomic dataset, the photoreceptor subpopulation was estimated to be significantly greater in the GP3 alpha than GP3 gamma MYC-associated subtype, and consequently is associated with a comparatively favorable clinical outcome. The presence of photoreceptor subpopulation in a subset of GP4 samples runs counter to the prior understanding of photoreceptor differentiation as a hallmark of GP3 and serves as an example of the underlying biological commonalities between GP3 and GP4 that confound subgroup assignment based on bulk-tumor sample analysis.

We show that neuronally-differentiated subpopulation SHH-C2 is the major cell type that constitutes SHH nodules. The SHH-C2 subpopulation was estimated to be the most abundant in infant subtypes SHH beta and gamma, further supporting the association of nodule formation with early neurodevelopment. SHH-C2 is particular abundant in the SHH gamma subtype that is largely composed of the MBEN histological type that carries a particularly favorable prognosis. Functional analyses of the molecular pathways driving differentiation toward the SHH-C2 is feasible now that this subpopulation has been molecularly defined and may reveal therapeutically relevant insights. This approach can explored by application of scRNA-seq to *in vivo* GEMM models, that we and others have shown to recapitulate the cellular heterogeneity of SHH MB (40).

The spectrum of MB infiltrating myeloid subpopulations revealed by our study is significantly greater than what was previously understood using flow and histological studies. A number of these myeloid subpopulations are distinguished either by neurodevelopment or antigen-presenting gene signatures. During early development, myeloid cells guide neural development, in part by interacting with developing neurons, phagocytosing apoptotic cells, pruning synapses, modulating neurogenesis, and regulating synapse plasticity and myelin formation (41). Two MB myeloid subpopulations exhibited neurodevelopmental-related characteristics, which differs from the immune role that has been assumed for MB tumor infiltrating immune cells. In SHH we identified an M2-myeloid subpopulation that was particularly abundant in nodular linings. In GP3 and GP4, we identified a complement-expressing myeloid subpopulation with potential neurodevelopmental roles that was associated with survival. These neurodevelopmentally-related myeloid subpopulations may be either subgroup or age restricted, M2-myeloid cells being specific to SHH that is predominant in infants. We identified a number of subpopulations harboring gene expression profiles indicative of active immune roles, in particular chemokine- and DC-like myeloid subpopulations that are both characterized by MHC class-II expression. Conversely, the presence of naïve microglia is consistent with previous observations of a particularly immunosuppressed microenvironment in MB (33). As this “cold” immunophenotype is likely to impede immunotherapy in MB, strategies targeting the specific immunophenotype of MB are necessary and can be advanced based on the findings of the present study.

Collectively, our study provides further insight into the neoplastic and immune cellular heterogeneity of the most common medulloblastoma subgroups and how this intra-tumoral heterogeneity is reflected in GEMM models of MB. These data show that tumor cells present a range of differentiation states that varies with each subgroup, and also show subgroup-specific interaction of the tumors with their immune microenvironments. We also provide interactive browsers for each of the described datasets that will facilitate the on-going interpretation of complex childhood MB biological data.

## Materials and methods

### Patient sample tumor cell preparation

MB samples for the scRNA-seq study were dissociated and viably banked at the time of surgery at our institution over a 10-year period. Patient material was collected at the time of surgery with consent (COMIRB 95-500). Samples were collected into PBS for subsequent scRNA-seq analysis and snap frozen for bulk-tumor methylome analysis (28 primary and 1 matched recurrence sample) (Supplementary Table 1). ScRNA-seq samples were rapidly dissociated into single cells using a mechanical process as described previously (33), viably cryopreserved and banked at <80°C for later use. In this way we were able to batch samples and thus limit experimental variance without compromising sample quality, as DMSO cryopreservation has been shown to match and in some cases exceed the quality of fresh samples with respect to scRNA-seq analysis (42).

### Methods for MP and MG GP3 MB GEMM models Tumor Generation and Tumor Cell Preparation

Cerebellar stem/progenitor cells (Prominin1+ cells) were purified by fluorescence activated cell sorting (FACS) from the cerebella of postnatal day 5–7 (P5–P7) C57BL/6J pups as previously described (34, 36). To generate MP or MG tumors, cells were infected with viruses produced from pMSCV-MycT58A-IRES-Luc and pMSCV-DNp53-IRES-GFP (for MP tumors) or pMSCV-GFI1-IRES-GFP (for MG tumors). After overnight infection, cells were washed and 10,000 cells were stereotactically injected into the cerebellum of 6-to 8-week-old NOD-SCID-IL2R-gamma (NSG) mice. Animals were monitored weekly and euthanized when they showed signs of tumor. Tumors were then enzymatically dissociated and viably cryopreserved prior to scRNA-seq analysis.

### Method for Math1-Cre/SmoM2 SHH GEMM tumor generation and tumor cell preparation

To generate mice with SHH-driven MB, we crossed SmoM2 mice (Jackson Labs, stock # 005130) with Math1-Cre (Jackson Labs, stock #011104) to generate M-Smo mice. All mice were of species Mus musculus and crossed into the C57BL/6 background through at least five generations. All animal studies were carried out with the approval of the University of North Carolina Institutional Animal Care and Use Committee under protocols (19-098). Mice were raised until P15 and then tumors were harvested under general anesthesia. Tumor samples were dissociated using the Papain Dissociation System (Worthington Biochemical) as previous described (18). Briefly, tumor samples were incubated in papain at 37°C for 15 min, then triturated and the suspended cells were spun through a density gradient of ovomucoid inhibitor. Pelleted cells were then cryopreserved prior to scRNA-seq analysis.

### Single-cell capture, RNA library preparation and sequencing

ScRNA-seq was performed as previously described (7). Samples were thawed in batches and flow sorted (Astrios EQ) to obtain viable single cells based on propidium iodide (PI) exclusion. With the study goal of performing scRNA-seq on 2,000 cells per sample, we utilized a Chromium Controller in combination with Chromium Single Cell V2 and V3 Chemistry Library Kits, Gel Bead & Multiplex Kit and Chip Kit (10X Genomics). Transcripts were converted to cDNA, barcoded and libraries were sequenced on Illumina NovaSeq6000 to obtain approximately 50 thousand reads per cell.

### ScRNA-seq data analysis

Raw sequencing reads were demultiplexed, mapped to the human reference genome (build GRCh38) and gene-expression matrices were generated using CellRanger (version 3.0.1). The resulting count matrices were further filtered in Seurat 3.1.0 (https://satijalab.org/seurat/) to remove cell barcodes with less than 200 genes or 500 UMIs, more than 30% of UMIs derived from mitochondrial genes, or more than 40,000 UMIs. This filtering resulted in 39,946 single cells across all samples (Supplementary Fig. 1). After normalization, these cells were clustered using the Seurat workflow based on dimensionality reduction by PCA using the 4,000 most variable genes. Coarse cell types were defined (immune, myeloid, and stroma cells) were assigned based on marker gene expression. Tumor cells were readily identified based on their discrete clustering patterns, and the presence of CNVs generated using inferCNV(v.1.3.6). Tumor cells from each subgroup were then reanalyzed separately. We applied Harmony alignment (theta = 1.5) to correct for inter-sample variation due to experimental or sequencing batch effects (8). After assessment of clustering using a variety of dimensions, we used 50 harmony dimensions to cluster the data and perform dimensionality reduction using Uniform Manifold Approximation and Projection (UMAP). Differential expression and marker gene identification was performed using Presto (43).

Chromosomal CNVs of single cells from scRNA-seq were inferred on the basis of average relative expression in variable genomic windows using inferCNV (https://github.com/broadinstitute/inferCNV). Cells classified as non-neoplastic were used to define a baseline of normal karyotype such that their average copy number value was subtracted from all cells.

Neoplastic subpopulations were characterized by direct examination of neoplastic-subpopulation specific gene lists, GO-term enrichment analysis, and inference of transcription factor regulatory networks. GO-term enrichments were calculated by gProfiler2 (v.0.1.9). Module scores for each subpopulation gene signature were generated using the top 200 markers from each subpopulation ranked by adjusted p-value, then secondarily ranked by positive fold change. Modules were calculated based on the methods in Tirosh *et al.*, as implemented in the Seurat function AddModuleScore (44). Single-cell regulatory network inference and clustering (pySCENIC) was used to identify transcription factor regulatory networks at the single cell level (9). Bipartite graphs were generated, using iGraph, between subpopulations using the correlation of normalized gene expression of shared variable genes as edge weights (39). Edges were removed if adjusted p-values were < 0.05 or if the correlations were less than 0.2 (GEMM to human comparisons) or less than 80% of the maximum correlations for human to human comparisons. One-to-one orthologs were used when comparing human and mouse single cell datasets and gene signatures. Harmony alignment (theta = 2) was performed to generate a UMAP projection containing mouse and human neoplastic subpopulations.

### Bulk-tumor methylome analyses

DNA was extracted from snap frozen EPN-PF surgical tumor samples (Qiagen, Allprep DNA/RNA mini kit). Twenty-eight surgical tumor samples were from initial presentation and 1 matched sample from a metastatic first recurrence. Methylome analysis of DNA from presentation samples was performed using the Illumina 850K methylation array. Resulting IDAT files were uploaded to MolecularNeuropathology.org (https://www.molecularneuropathology.org/mnp) which provided subgrouping into MB molecular subgroups/subtypes, and chromosomal copy number variants (CNVs) (Supplementary Table 1).

### Deconvolution

Cibersort was used to perform deconvolution of bulk-tumor MB transcriptomes using scRNA-seq subpopulation signature genes (45). The combined MAGIC transcriptomic dataset (n=747) was unlogged and used as the mixture file. A signature gene input matrix was generated from log2 values of normalized raw scRNA-seq expression data averaged for neoplastic and immune clusters (Supplementary Table 2). Cibersort was run using these datasets with 100 permutations.

### Immunohistochemistry

Immunohistochemistry was performed on 5-μm formalin-fixed, paraffin-embedded (FFPE) tumor tissue sections using a Ventana autostainer. Antigen retrieval was performed by incubation in citrate solution pH 6.0 for 10 min at 110°C. Slides were treated with primary antibodies for GRIA2 (Abcam (clone EP929Y), 1:200 dilution), IMPG2 (Novus (Cat# NBP2-58919), 1:400), MRC1 (Sigma (clone 5C11), 1:500) and STMN2 (Novus (Cat#NBP1-49461), 1:1000) for 32 minutes at 37°C. All immunostained sections were counterstained with hematoxylin. Neuropathological review of staining and blinded IHC scoring was then performed (N.W., A.M.D.).

### Survival studies

Survival data was obtained for a subset of the MAGIC transcriptomic cohort (n=539). Survival analyses were performed using Prism (GraphPad) software, with outcome censored at 5 years. Hazard ratios (HRs) for progression free survival (PFS) and overall survival (OS) were estimated using log-rank (Mantel–Cox) analysis of high versus low (i) expression of each subpopulation as defined by median deconvoluted subpopulation fractions, and (ii) IHC as defined by median score or present versus absent.

### Data and Code Availability

Custom code used in this study is available at https://github.com/rnabioco/medulloblast. ScRNAseq and methylation data have been deposited in the National Center for Biotechnology Information Gene Expression Omnibus (GEO) database and are publicly accessible through GEO SuperSeries accession number GSE156053 (https://www.ncbi.nlm.nih.gov/geo/query/acc.cgi?acc=GSE156053). All MB scRNAseq data is also available as a browsable webresource at the Pediatric Neuro-oncology Cell Atlas (pneuroonccellatlas.org).

### Statistics

Statistical analyses were performed using R, Prism (GraphPad), and Excel (Microsoft) software. Details of statistical tests performed are included in Fig. legends. For all tests, statistical significance was defined as P < 0.05.

## Author Contributions

A.M.D. and S.V., performed experiments with assistance from A.N., A.M.G., V.A., A.G., T.C.H and M.H.H. K.A.R. performed bioinformatic analyses and B.S., S.J.W., D.S.M. and M.S. made additional bioinformatic contributions. N.W. assisted with neuropathology concerns. T.R.G., A.G., R.J.W-R. and J.O. provided GEMM samples. K.A.R., R.V., S.M., J.R.H., N.K.F. and A.M.D. were responsible for the design of the study. All authors assisted with manuscript preparation.

## Acknowledgements

K.A.R. is supported as an Informatics Fellow of the RNA Bioscience Initiative, University of Colorado School of Medicine. This study was supported by the Morgan Adams Foundation. The University of Colorado Denver Genomics and Microarray, Flow Cytometry, and Histology Shared Resources are supported by the University of Colorado’s NIH/NCI Cancer Center (P30CA046934).

## Supplementary Tables

**Supplementary Table 1.** Patient and sample details.

**Supplementary Table 2.** Cibersort neoplastic and immune subpopulation signatures.

**Supplementary Table 3.** Neoplastic subpopulation marker genes.

**Supplementary Table 4**. Neoplastic subpopulation GOTerm enrichments.

**Supplementary Table 5.** Neoplastic subpopulation inferred transcription factor activity.

**Supplementary Table 6.** Survival analyses of deconvoluted MAGIC cohort subpopulation proportions.

**Supplementary Table 7.** Correlation matrix for deconvoluted MAGIC cohort subpopulation proportions.

**Supplementary Table 8.** Immune subpopulation marker genes.

**Supplementary Table 9.** GEMM subpopulation marker genes.

