## Supplemental-Figures for "Neoplastic and immune single cell transcriptomics define subgroup-specific intra-tumoral heterogeneity of childhood medulloblastoma"

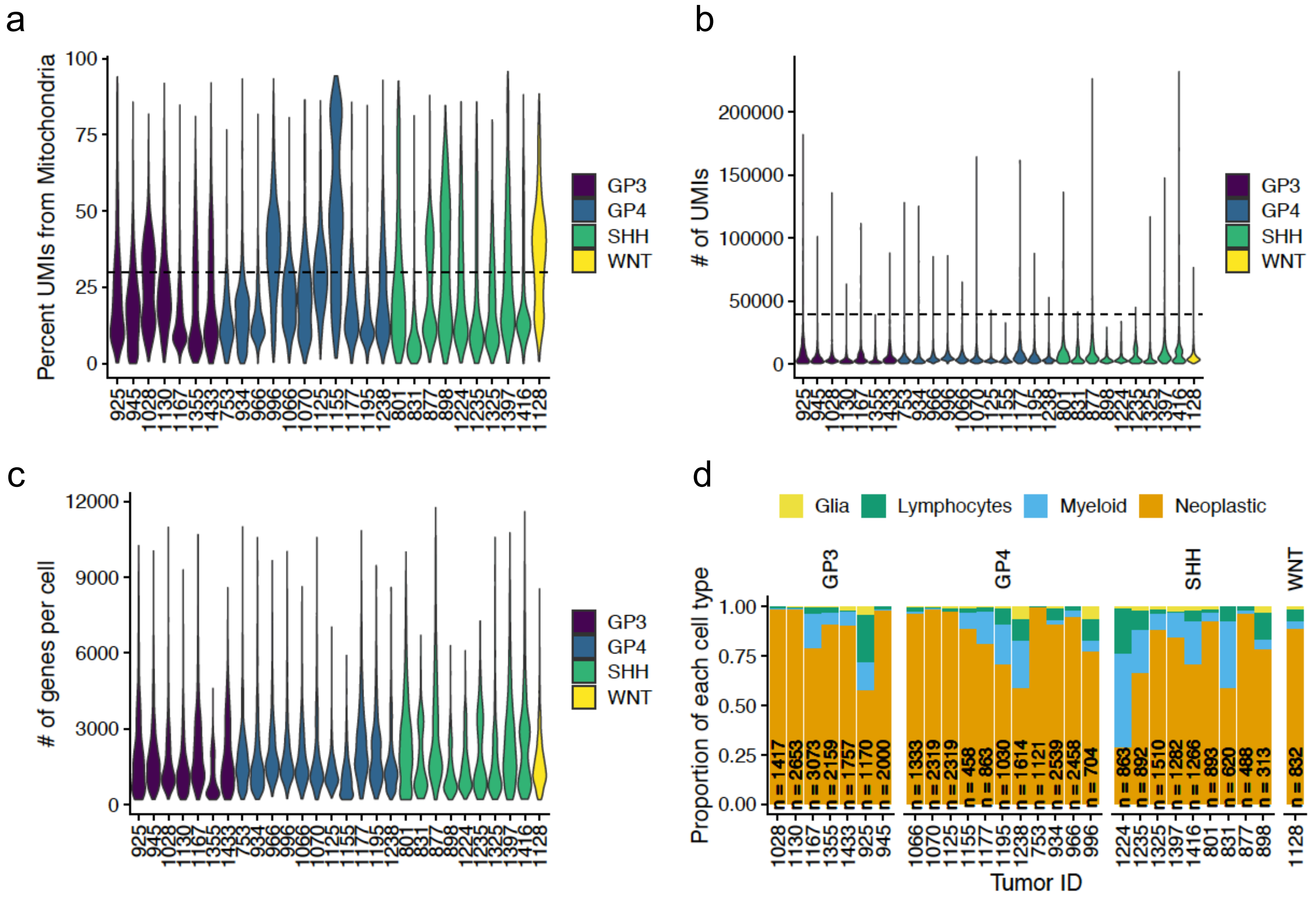


**Supplementary Fig. 1. QC metrics on 29 MB patient samples.** **a** Percent mitochondrial reads, **b** number of unique molecular identifiers (UMI) and **c** number of genes per cell. The pre-filtering plots have dotted lines (thresholds) to show how filtering was performed. **d** Proportion of cells from each sample grossly assigned to neoplastic and non-neoplastic cell types.

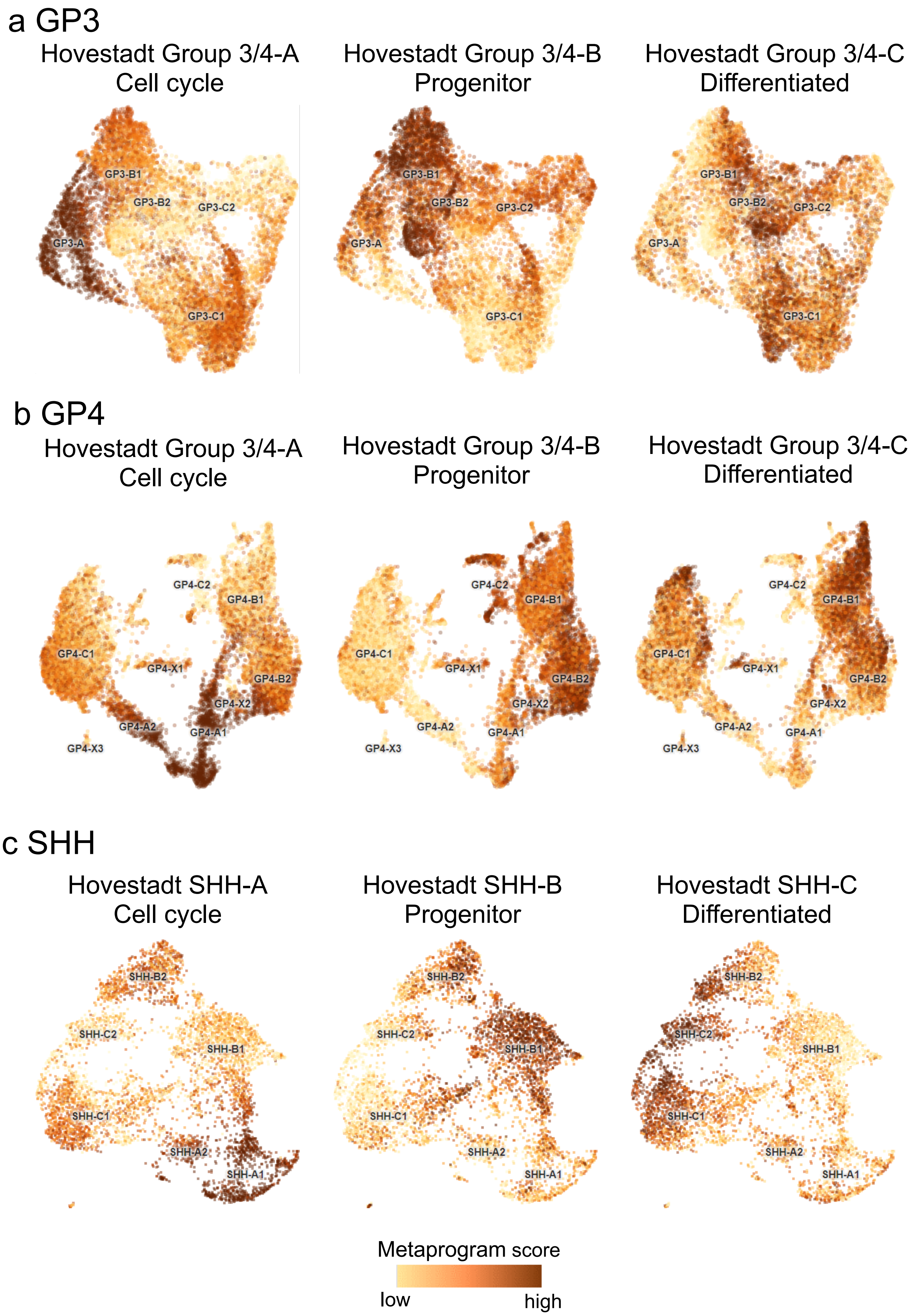
a GP3
Hovestadt Group 3/4-A
Cell cycle
Hovestadt Group 3/4-B
Progenitor
Hovestadt Group 3/4-C
Differentiated
GP3-B1
GP3-B2
GP3-C2
GP3-A
GP3-C1
b GP4
Hovestadt Group 3/4-A
Cell cycle
Hovestadt Group 3/4-B
Progenitor
Hovestadt Group 3/4-C
Differentiated
GP4-C2
GP4-B1
GP4-C1
GP4-X1
GP4-B2
GP4-X2
GP4-A2
GP4-A1
GP4-X3
c SHH
Hovestadt SHH-A
Cell cycle
Hovestadt SHH-B
Progenitor
Hovestadt SHH-C
Differentiated
SHH-B2
SHH-C2
SHH-B1
SHH-C1
SHH-A2
SHH-A1
Metaprogram score
low
high

**Supplementary Fig. 2. MB metaprogram scores in MB subgroup subpopulations.** Harmony aligned **a** GP3, **b** GP4 and **c** SHH neoplastic cells colored according to enrichment of corresponding A – cell cycle, B – progenitor and C – differentiated transcriptional metaprograms, as described by Hovestadt *et al.* (5).

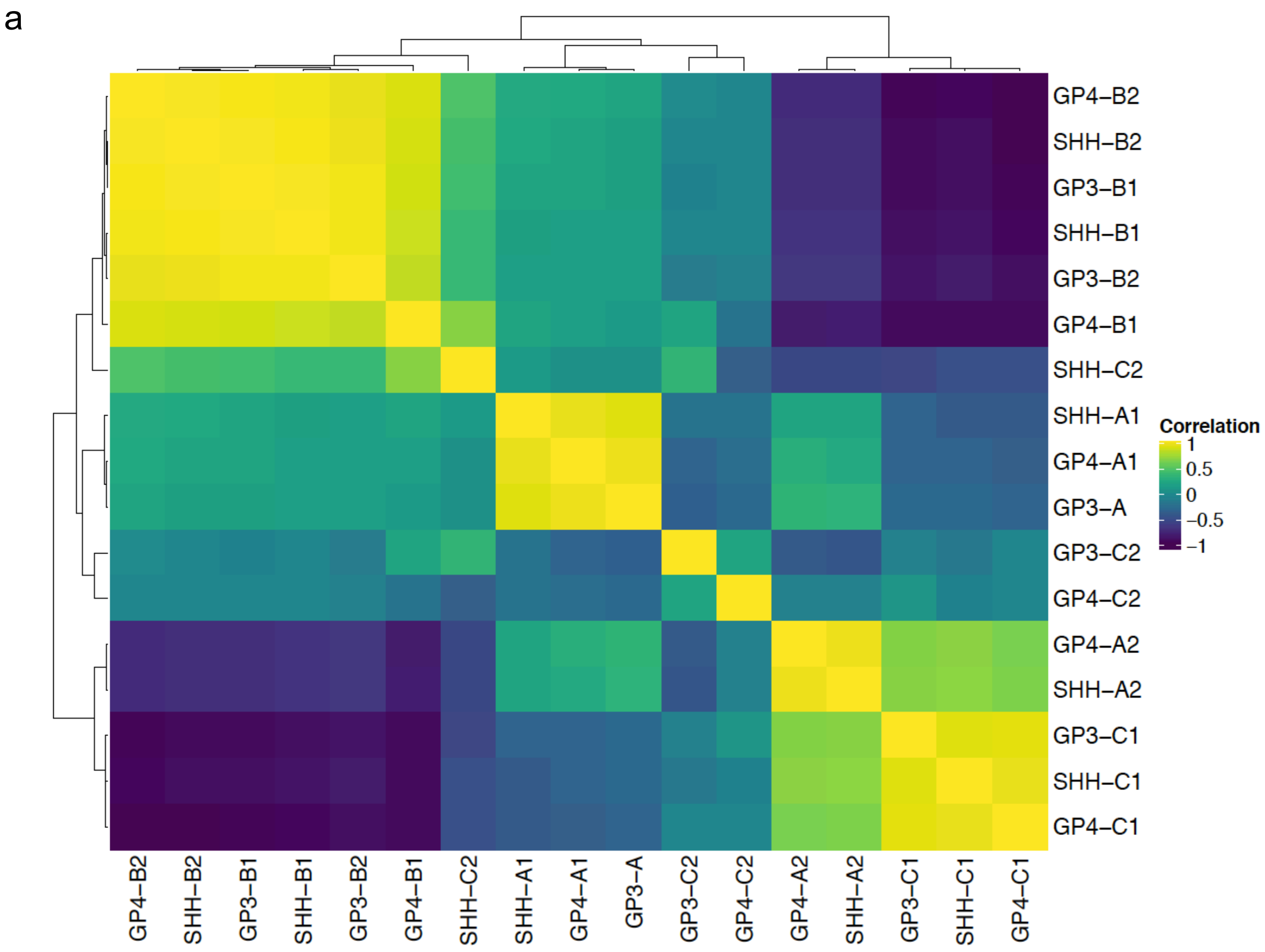


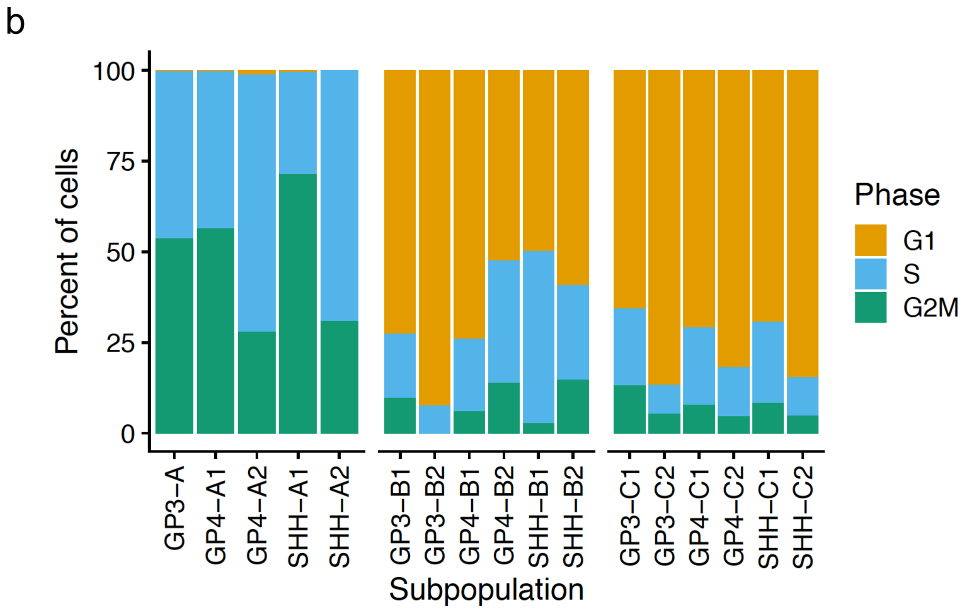


**Supplementary Fig. 3. a** Correlation matrix of tumor subpopulation scores for all tumor cells. For each tumor subpopulation the top 200 markers are used to generate a subpopulation score for all tumor cells. Every cell then has a subpopulation score from all of the tumor subpopulations. Spearman correlation coefficients then are calculated between each subpopulation score. **b** Proportion of cells in G1, S and G2M cell cycle phases in MB neoplastic subpopulations.

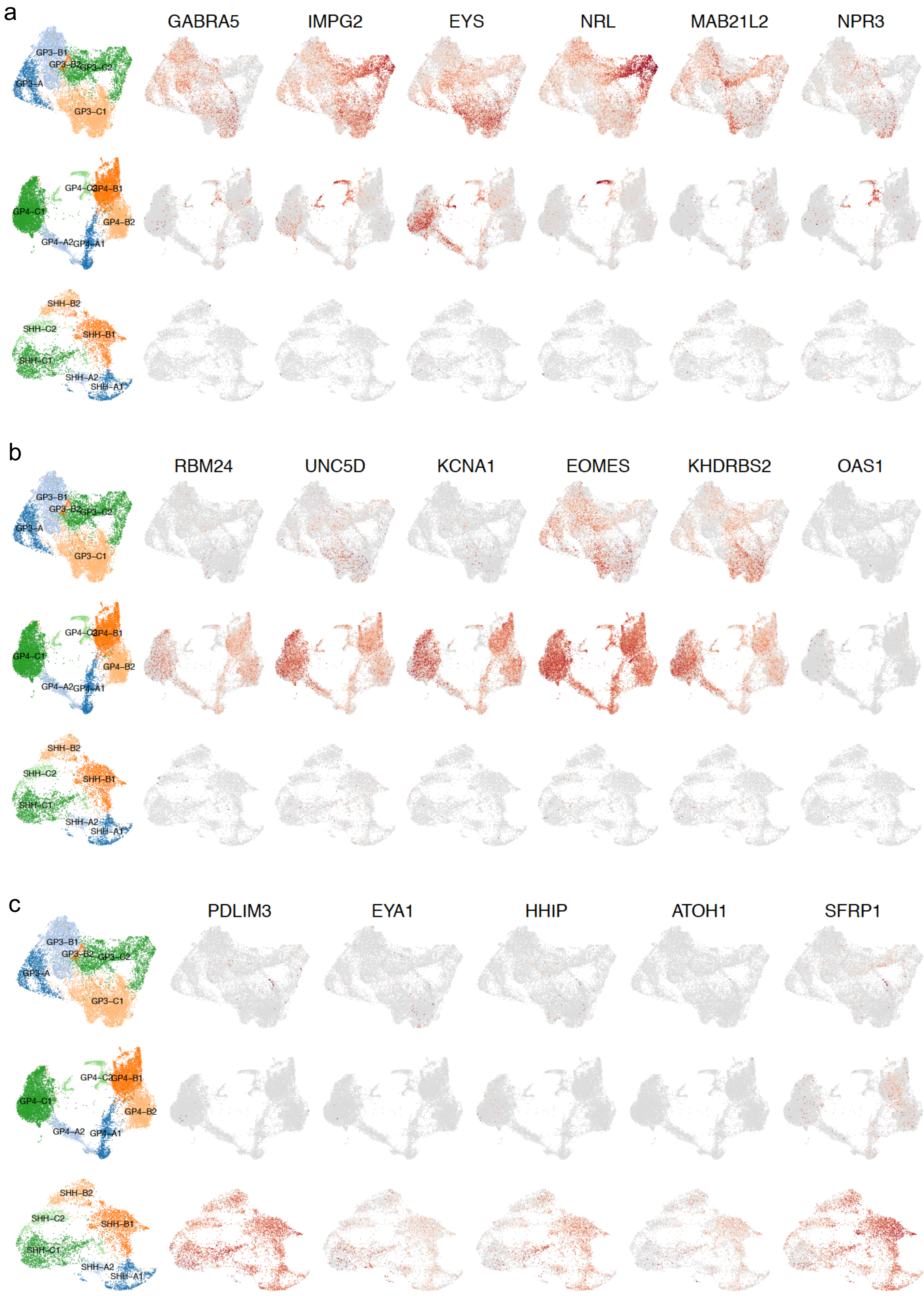
a
GABRA5
IMPG2
EYS
NRL
MAB21L2
NPR3
GP3–B1
GP3–B2
GP3–C2
GP3–A
GP3–C1
GP4–C2
GP4–B1
GP4–C1
GP4–B2
GP4–A2
GP4–A1
SHH–B2
SHH–C2
SHH–B1
SHH–C1
SHH–A2
SHH–A1
b
RBM24
UNC5D
KCNA1
EOMES
KHDRBS2
OAS1
c
PDLIM3
EYA1
HHIP
ATOH1
SFRP1

**Supplementary Fig. 4. Subpopulation distribution of consensus MB subgroup markers.** UMAP feature plots depicting expression of consensus MB subgroup nanostring markers (14) for **a** GP3, **b** GP4 and **c** SHH neoplastic subpopulations. Expression is indicated by color scale for each gene. Key indicates location of major MB neoplastic subpopulations for each subgroup.

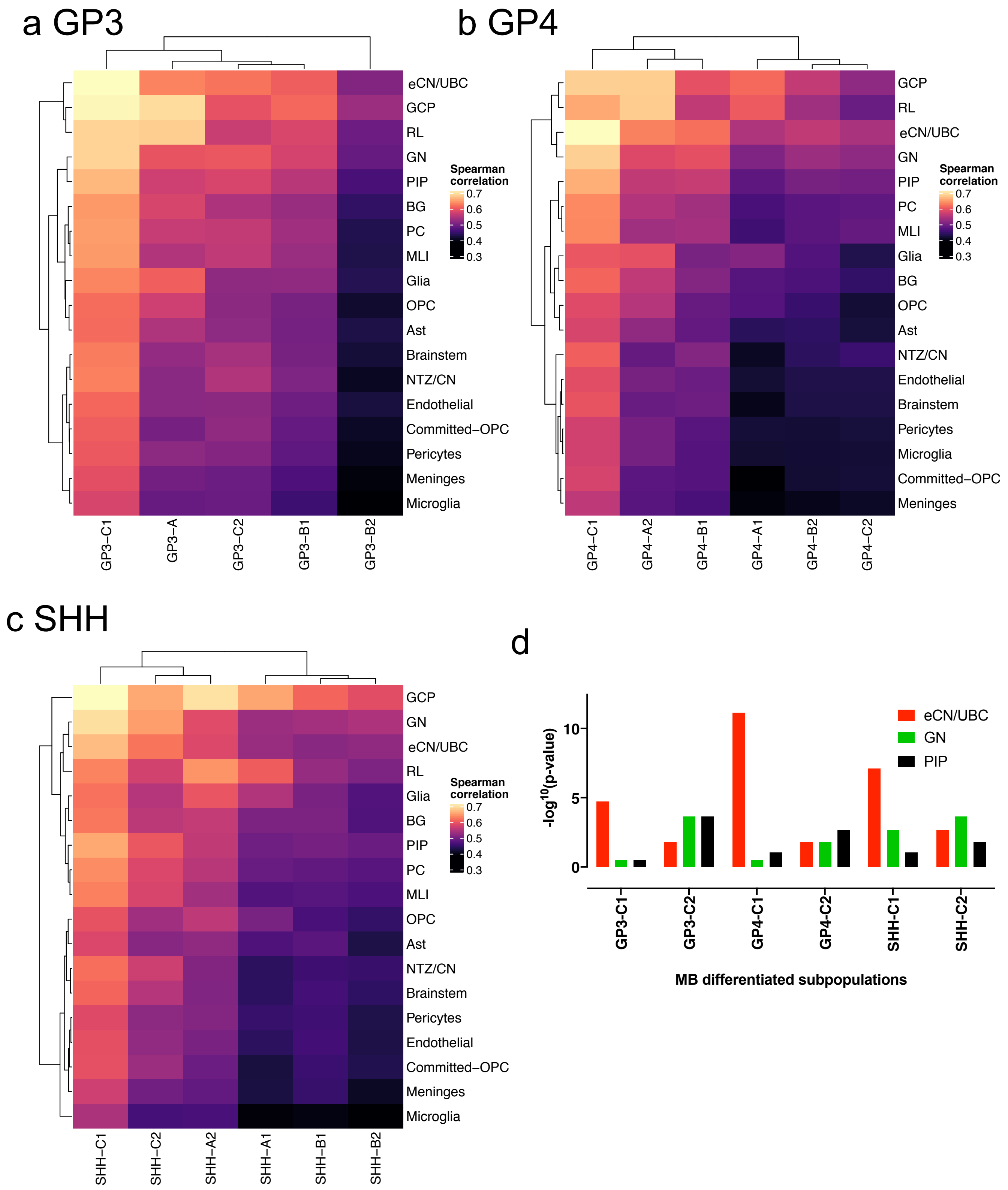


**Supplementary Fig. 5. Human cerebellar developmental cell lineage signature enrichment MB neoplastic subpopulations.** Heatmaps depicting correlation (Spearman's R) of gene expression from scRNA-seq analysis of human cerebellar developmental cell lineages (17) compared to MB **a** GP3, **b** GP4 and **c** SHH neoplastic subpopulations. **d** Top 3 enriched scRNA-seq cerebellar lineages signatures in MB neoplastic differentiated subpopulations (hypergeometric test p-value). Abbreviations: Ast, astrocyte; BG, basal ganglia; CN, cerebellar nuclei; eCN, excitatory CN; GCP, granule cell precursor; GN, granule neurons; NTZ, nuclear transitory zone; MLI, molecular layer interneurons; OPC, oligodendrocyte progenitor cell; PC, purkinje cell; PIP, PAX2+ interneuron progenitors; UBC, unipolar brush cell.

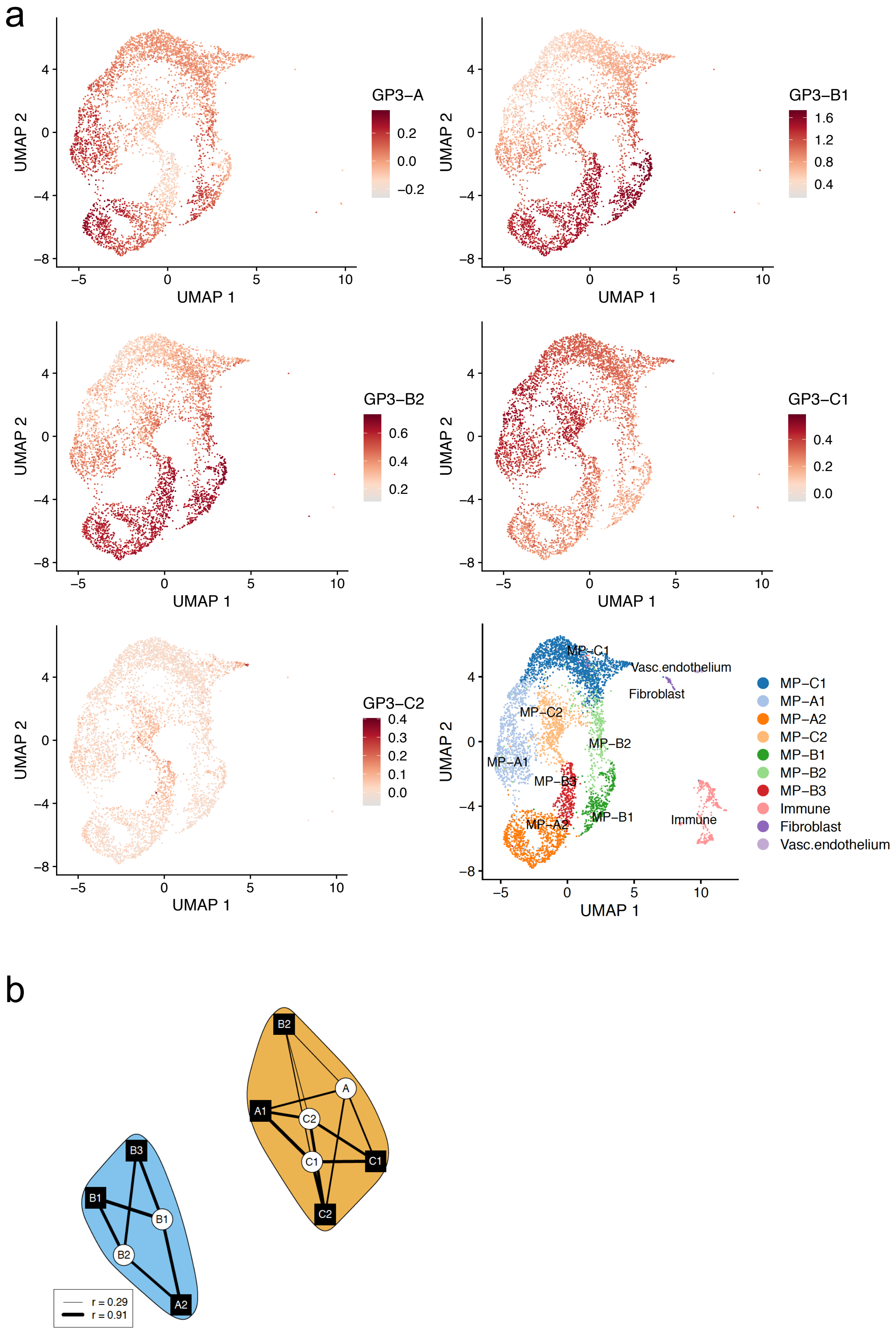


**Supplementary Fig. 6. Cellular distribution of human MB subpopulation signatures in GP3 model MP. a** UMAP feature plots depicting expression of human MB neoplastic subpopulation metasignatures (brown) for MYC-DNp53 driven GP3 GEMM (MP). Key indicates location of major MB neoplastic subpopulations for each subgroup. **b** Directed bi-partite graph comparing GEMM with human neoplastic subpopulations from the corresponding subgroup.

a

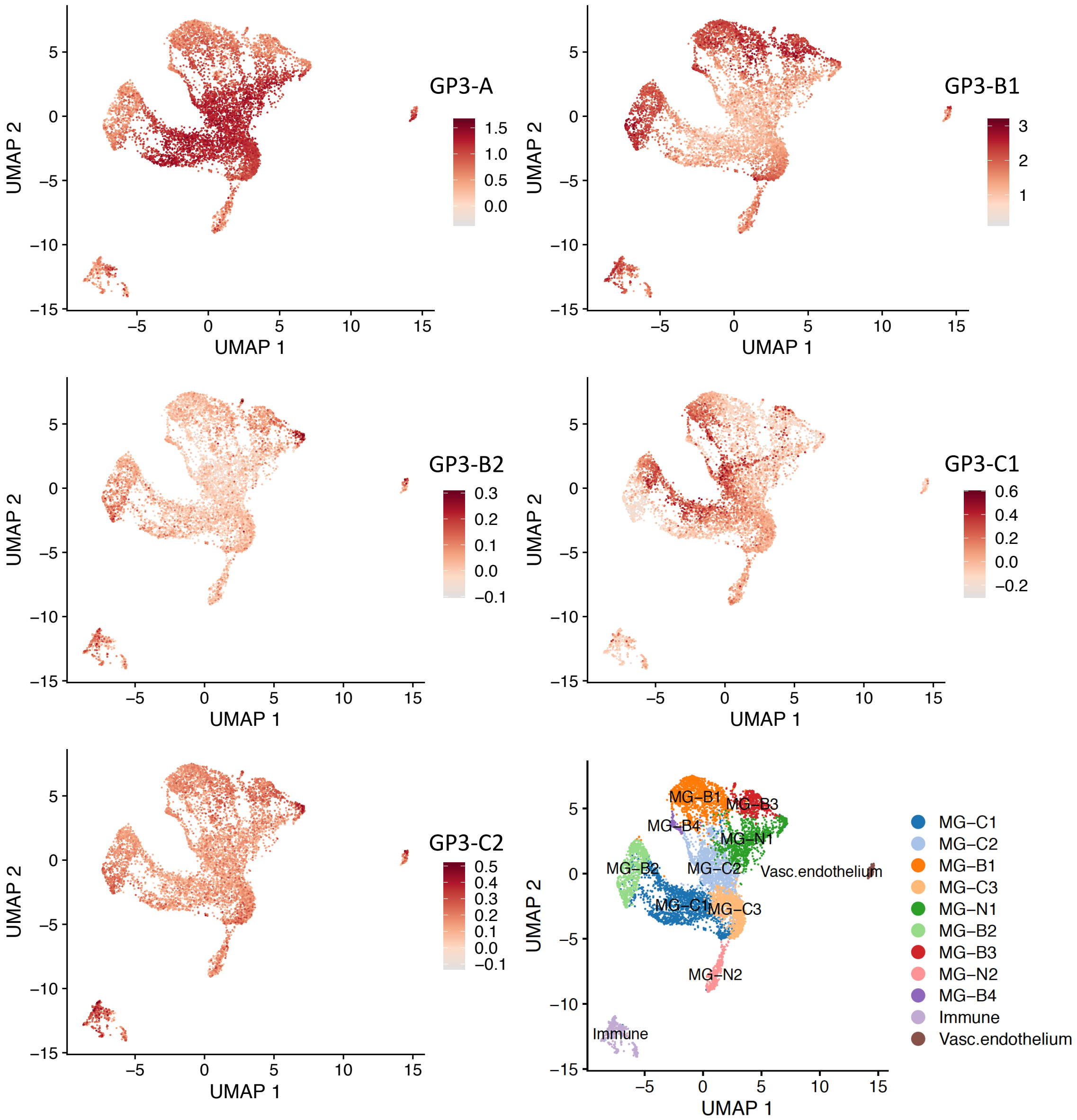


b

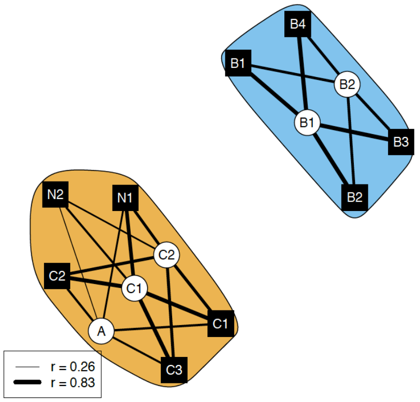


**Supplementary Fig. 7. Cellular distribution of human MB subpopulation signatures in GP3 model MG. a** UMAP feature plots depicting expression of human MB neoplastic subpopulation metasignatures (brown) for MYC-GFI1 driven GP3 GEMM (MG). Key indicates location of major MB neoplastic subpopulations for each subgroup. **b** Directed bi-partite graph comparing GEMM with human neoplastic subpopulations from the corresponding subgroup.

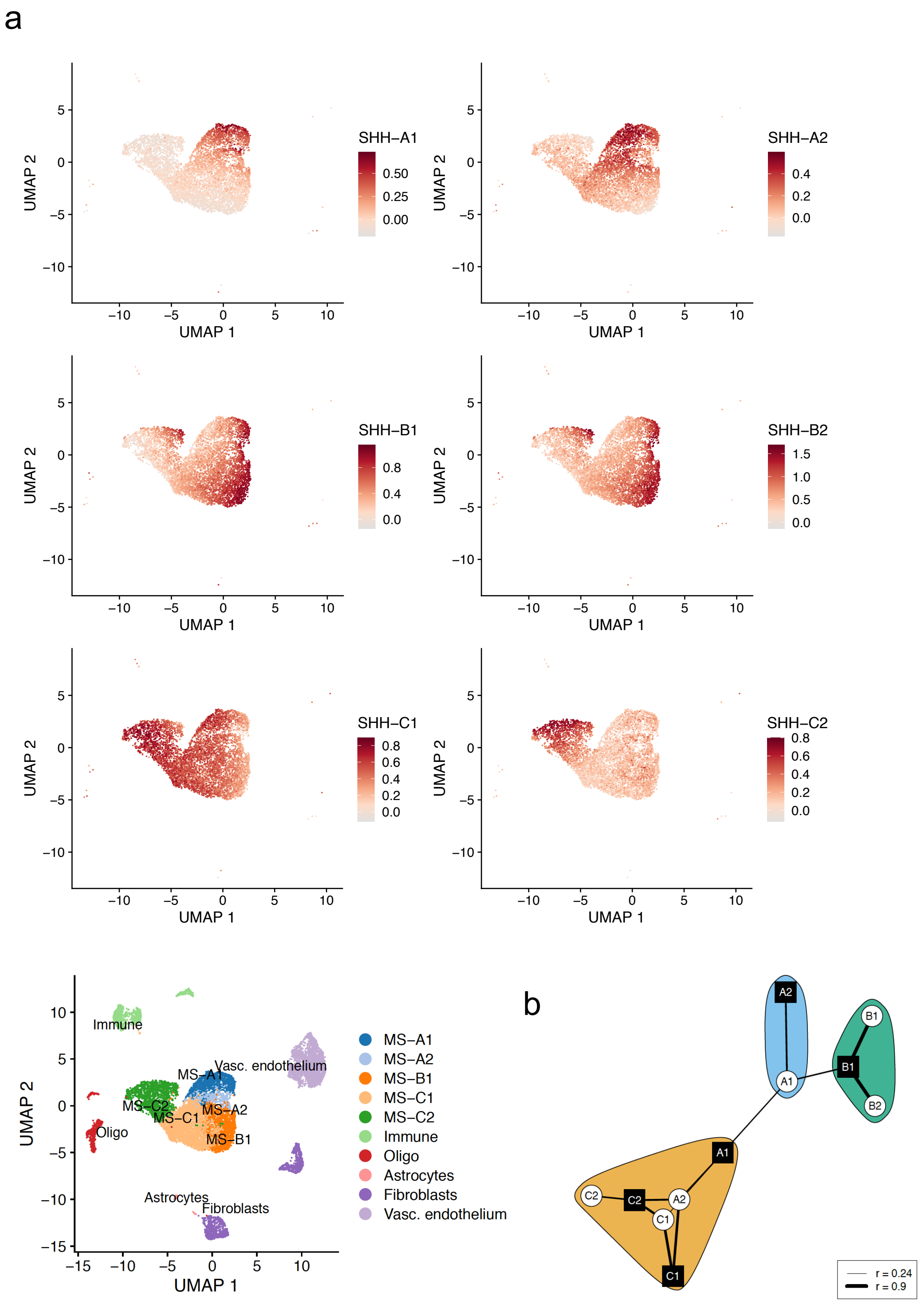


**Supplementary Fig. 8. Cellular distribution of human MB subpopulation signatures in SHH model MS. a** UMAP feature plots depicting expression of human MB neoplastic subpopulation metasignatures (brown) for Math1/SmoM2 driven SHH GEMM (MS). Key indicates location of major MB neoplastic subpopulations for each subgroup. **b** Directed bi-partite graph comparing GEMM with human neoplastic subpopulations from the corresponding subgroup.
